# Relaxing parametric assumptions for non-linear Mendelian randomization using a doubly-ranked stratification method

**DOI:** 10.1101/2022.06.28.497930

**Authors:** Haodong Tian, Amy M. Mason, Cunhao Liu, Stephen Burgess

## Abstract

Non-linear Mendelian randomization is an extension to standard Mendelian randomization to explore the shape of the causal relationship between an exposure and outcome using an instrumental variable. The approach divides the population into strata and calculates separate instrumental variable estimates in each stratum. However, the standard implementation of stratification in non-linear Mendelian randomization, referred to as the residual method, relies on strong parametric assumptions of linearity and homogeneity between the instrument and the exposure to form the strata. If these stratification assumptions are violated, the instrumental variable assumptions may be violated in the strata even if they are satisfied in the population, resulting in misleading estimates. We propose a new stratification method, referred to as the doubly-ranked method, that does not require strict parametric assumptions to create strata with different average levels of the exposure such that the instrumental variable assumptions are satisfied within the strata. Our simulation study indicates that the doubly-ranked method can obtain unbiased stratum-specific estimates and appropriate coverage rates even when the effect of the instrument on the exposure is non-linear or heterogeneous. Moreover, it can also provide unbiased estimates when the exposure is coarsened (that is, rounded, binned into categories, or truncated), a scenario that is common in applied practice and leads to substantial bias in the residual method. We applied the proposed doubly-ranked method to investigate the effect of alcohol intake on systolic blood pressure, and found evidence of a positive effect of alcohol intake, particularly at higher levels of alcohol consumption.

## Introduction

Mendelian randomization is an epidemiological technique that uses genetic variants as instrumental variables to make causal inferences from observational data^1;2^. An extension to the method, known as non-linear Mendelian randomization, first divides the population into strata with different average levels of the exposure, and then performs separate instrumental variable analyses in each stratum to obtain stratum-specific estimates, referred to as localized average causal effect (LACE) estimates^3;4^. This allows researchers to explore the shape of the causal relationship between the exposure and the outcome. While other methods have been proposed for performing non-linear instrumental variable analysis^5–7^, such methods typically require the parametric model relating the exposure to the outcome to be specified, or else perform model selection amongst a set of models^8^. Inferences from such approaches can be sensitive to the specification of the non-linear function or model selection procedure^9^. Additionally, it is difficult to fit detailed non-linear models if the instrumental variable takes a small number of discrete values, or explains a small proportion of the variance in the exposure; both of these scenarios are common in Mendelian randomization.

The stratification method for non-linear Mendelian randomization has been used to investigate the shape of the exposure–outcome relationship for body mass index (BMI) with mortality^10^, systolic blood pressure with cardiovascular disease^11^, and vitamin D with a range of outcomes^12^. For vitamin D, instrumental variable estimates for all-cause mortality were null in overall analyses and for strata of the population with adequate (*>*50 nmol/L) levels of 25-hydroxyvitamin D [25(OH)D], a metabolite of vitamin D used as a clinical indicator of vitamin D status, but in the protective direction for strata of the population with deficient (*<*25 nmol/L) and insufficient (25−50 nmol/L) levels of 25(OH)D. This suggests that vitamin D supplementation has a threshold effect on all-cause mortality risk, indicating that supplementation may only be beneficial for all-cause mortality for those with low vitamin D status.

An important technical detail in these analyses is that stratification is not performed on the exposure directly. The reason is that the exposure is a collider in the standard directed acyclic graph for an instrument variable: it is a common effect of the instrument and the exposure–outcome confounders (Figure 1)^13^. If we regard the instrumental variable as equivalent to random allocation in a randomized trial, the exposure is a post-randomization covariate, and so stratification on the exposure is inappropriate^14^. The original proposal for stratified non-linear Mendelian randomization was to first calculate a variable referred to as the “residual exposure” and stratify on this^3^. The residual exposure is calculated by regressing the exposure on the genetic instrument (either a single genetic variant or a score comprising multiple genetic variants), and taking the residual from this equation. By the properties of linear regression, the residual exposure is independent of the genetic instrument, and so strata defined using the residual exposure are independent of the genetic instrument. A weakness of this approach is that it relies on strong parametric assumptions of linearity and no effect modification between the genetic instrument and the exposure^15^.

**Figure 1:**
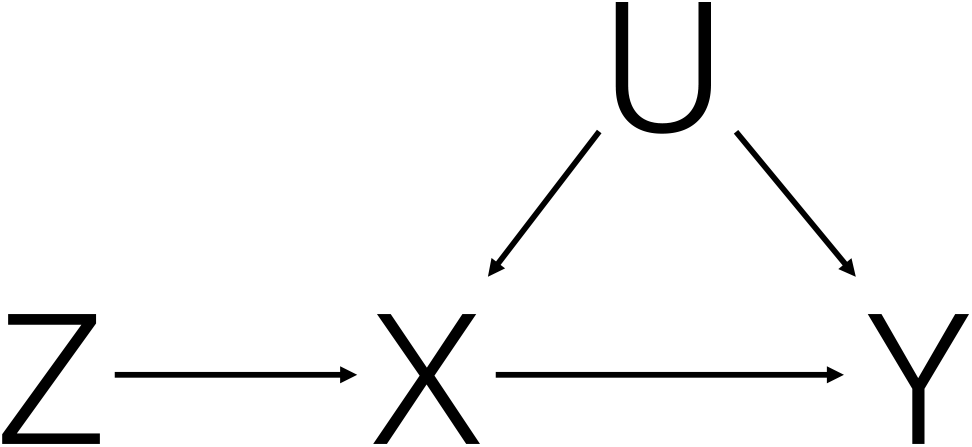
Directed acyclic graph (DAG) illustrating the instrumental variable assumptions. The exposure is denoted as *X*, the genetic instrument as *Z*, the outcome as *Y*, and exposure–outcome confounders as *U*. The exposure *X* is a collider in this DAG, as it is a common effect of the instrument and confounders.

In this paper, we propose a new stratification method that does not require strict parametric assumptions to create strata of the population that have different average levels of the exposure, but are independent of the instrument. We demonstrate in a simulation study that the method, referred to as the doubly-ranked stratification method, can obtain unbiased LACE estimates when the effect of the instrument on the exposure is non-linear or heterogeneous. In contrast, LACE estimates from the residual stratification method are biased in these scenarios. We also consider a scenario in which the exposure is coarsened (that is, either rounded or binned into categories), as is the case for several epidemiological risk factors. This scenario represents a difficulty for the residual stratification method, but can be accommodated by the doubly-ranked method. We apply the doubly-ranked method to study the shape of the causal effect of alcohol intake on systolic blood pressure (SBP). Software for implementing the doubly-ranked method is available as part of the SUMnlmr^16^ package at https://github.com/amymariemason/SUMnlmr.

## Methods

### Assumptions and data set-up

We assume the existence of a genetic instrument that satisfies the core instrumental variable assumptions^17;18^:

i. the instrument is associated with the exposure (relevance);
ii. the instrument is not associated with the outcome via a confounding pathway (exchange-ability);
iii. the instrument does not affect the outcome directly, only possibly indirectly via the exposure (exclusion restriction).

For interpretability of the instrumental variable estimates, we additionally make either the monotonicity or homogeneity estimation assumption^19^. The monotonicity assumption states that the effect of the instrument on the exposure is increasing for all individuals in the population (or alternatively, it is decreasing for all individuals in the population). There are various versions of the homogeneity assumption; the simplest version states that the effect of the instrument on the exposure is constant for all individuals in the population^20^. Under the monotonicity assumption, the instrumental variable estimate represents a local average treatment effect; under the homogeneity assumption, it represents an average treatment effect. The LACE is defined as the average causal effect for a subset of the population defined by stratification^3^. Depending on the estimation assumption, it either represents a local average treatment effect (monotonicity) or an average treatment effect (homogeneity) for that subset of the population. For a single instrument, this estimate can be calculated using the ratio method: by dividing the genetic association with the outcome by the genetic association with the exposure^21^.

We note that even though the instrument is not associated with confounders in the overall population, it may be associated in a subset of the population defined by stratification. The other two core IV assumptions (relevance and exclusion restriction) should hold in any subset of the population.

We denote the genetic instrument as *Z*, the exposure as *X*, the outcome as *Y*, and confounders of the exposure–outcome association (assumed unmeasured) as *U*. We first introduce the residual and doubly-ranked stratification methods, and then explore their properties in a simulation study and an applied analysis.

### Residual stratification method

Under the stratification assumption that the effect of the genetic instrument on the exposure is linear and homogeneous, the residual from regression of the exposure on the instrument will represent the value of the exposure as if the genetic instrument took the value zero. This variable, known as the ‘residual exposure’, is typically highly correlated with the exposure, as genetic instruments typically do not explain a large proportion of variance in the exposure. However, if we considered stratifying on the exposure directly, the distribution of the instrument would be different in the various strata. Values of the genetic instrument corresponding to increased levels of the exposure would be more common in strata with greater levels of the exposure. In contrast, not only is the residual exposure uncorrelated with the instrument, such that the instrument should be distributed on average similarly in the different strata; but, under the linearity and homogeneity assumptions, the functional dependence of the residual exposure on the instrument is broken. As the exposure is a common effect of the genetic instrument and confounders, the genetic instrument and confounders will be correlated within strata of the exposure; this is an example of collider bias^13^. However, as the residual exposure is not an effect of the genetic instrument, the genetic instrument and confounders will remain uncorrelated within strata of the residual exposure. We can therefore obtain LACE estimates within strata of the residual exposure.

### Doubly-ranked stratification method

For simplicity of explanation, we initially assume that we have a sample size of 1000, and the instrumental variable takes 100 different values 10 times each: that is, 10 individuals have *Z* = 1, 10 individuals have *Z* = 2, and so on. We first rank individuals into 100 pre-strata according to their level of the instrument, such that all those in pre-stratum *j* have *Z* = *j*. Then, within each pre-stratum we rank individuals according to their level of the exposure, and divide into strata. The first stratum consists of the individual who has the lowest value of the exposure in pre-stratum 1, the individual who has the lowest value of the exposure in pre-stratum 2, the individual who has the lowest value of the exposure in pre-stratum 3, and so on. The second stratum consists of the individual who has the second lowest value of the exposure in pre-stratum 1, the individual who has the second lowest value of the exposure in pre-stratum 2, the individual who has the second lowest value of the exposure in pre-stratum 3, and so on. We end up with 10 strata, each of which contains one individual from pre-stratum 1, one individual from pre-stratum 2, one individual from pre-stratum 3, and so on up to pre-stratum 100. The method is named doubly-ranked due to the two stratifications based on successive rankings: first, stratification into pre-strata based on the levels of the instrument; and then stratification into final strata based on the levels of the exposure within each pre-stratum.

In this simplified example, the distribution of the instrument is identical in each stratum by design; all strata have one individual with each value of the instrument from 1 to 100. As the stratification variable is not correlated with the instrument, the instrument should remain uncorrelated with confounders within each stratum. By construction, the average level of the exposure is increasing across the different strata. Hence, LACE estimates can be obtained within these strata which inform the investigator about average treatment effects (or local average treatment effects) for subsets of the population with different average levels of the exposure.

Suppose now that the instrument takes different values for a sample size of *N* = *J* × *K*, and we want to divide the population into *J* strata each containing *K* individuals. The doubly-ranked method works similarly as before: first, we rank individuals based on their level of the instrument, and divide into *K* pre-strata each containing *J* individuals; then we rank individuals within each pre-stratum based on their level of the exposure. We put the individuals with the lowest level of the exposure from each pre-stratum into stratum 1, the individuals with the second lowest level of the exposure from each pre-stratum into stratum 2, and so on. In this situation, the distribution of the instrument will not be exactly identical within each of the strata, but they should be similar by construction. Provided the effect of the instrument on the exposure is not too strong, correlation between the instrument and stratification will be low. Moreover, there is no functional dependence of stratification on the instrument. So, for a valid instrument, we expect correlation between the instrument and the confounders within the strata to be negligibly low. If some individuals have exactly the same value of the instrument or exposure, we break ties at random.

As in the residual method, we can obtain LACE estimates within strata of the population. As the method allows the genetic effect on the exposure to vary, we calculate the genetic associations with the exposure (and outcome) separately in each stratum to obtain the LACE estimates. For comparability of the methods, here we also estimate the genetic associations with the exposure within each stratum for the residual stratification method; if the effect of the genetic variant on the exposure is truly homogeneous, then it could be estimated more precisely in the whole population. A diagram outlining the doubly-ranked method is provided as Figure 2.

**Figure 2:**
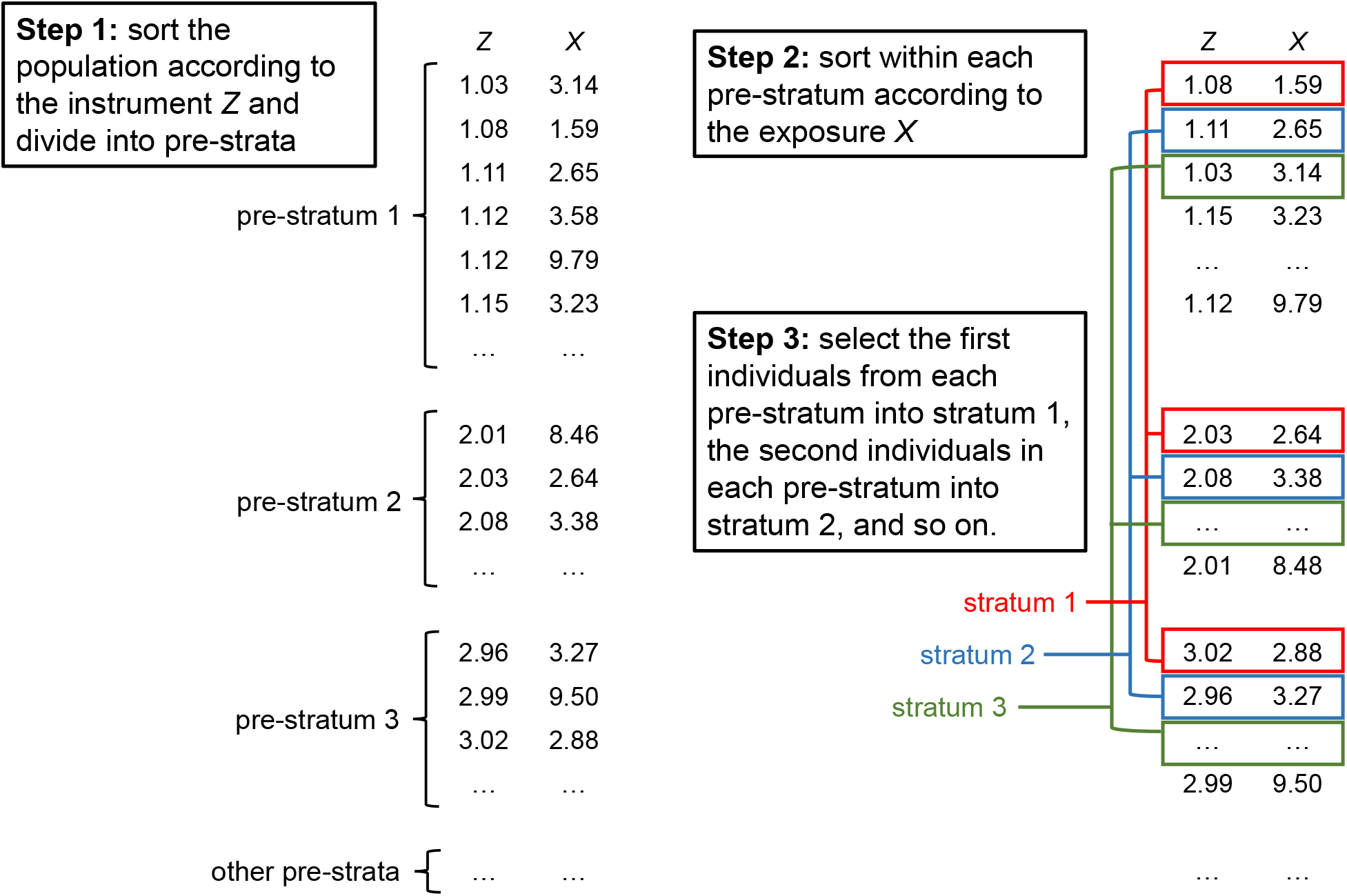
Schematic diagram illustrating the doubly-ranked stratification method.

### Coarsened exposure values

The residual stratification method requires values of the exposure to be known precisely, otherwise it is not possible to calculate the residual exposure in a way that is not causally dependent on the instrument values. However, in practice many exposure variables are coarsened^22^. For example, alcohol intake is often measured in categories (such as 0-5 g/day, 5-10 g/day, and so on), not absolute amounts; age at menarche is typically reported as a whole number of years; and manual blood pressure measurements are often preferentially reported as multiples of 5. This is a particular problem when the categorization is more coarse then the effect of the instruments; for example, sleep duration is often reported as a whole number of hours, but the average effect of genetic variants on sleep duration is far less than 1 hour. This means that boundaries between strata would either divide individuals according to their exposure values, introducing collider bias, or else they would divide individuals for a given value of the exposure according to their instrument values, creating strata with irregular distributions of the instrument.

In contrast, the doubly-ranked method should be able to form meaningful strata even if the exposure is coarsened. A further practical feature of the doubly-ranked method is that strata are equal in size even if the distribution of the exposure is irregular.

If the exposure is coarsened to take a small number of values, then it is not possible to divide the population into a large number of strata in a meaningful way. In an extreme case, it may be that all the individuals in a stratum have the same value of the exposure, in which case the relevance assumption does not hold. There is also a possible violation of the exchangeability assumption when coarsening of the exposure leads to patterns arising within the strata. We have developed a Gelman–Rubin uniformity statistic to assess whether clumps in the coarsened exposure distribution (that is, groups of individuals with the same coarsened exposure value) are distributed uniformly within the strata; high values of this statistic indicate potential violation of the exchangeability assumption, and suggest the use of fewer strata (see Text S1).

### Simulation study

We explore how the residual and doubly-ranked stratification methods perform in a range of simulation scenarios. We consider three causal relationships between the exposure and the outcome:

1. No causal effect of the exposure on the outcome: *Y* = *U* + *ϵ*_*Y*_.
2. U-shaped causal effect of the exposure on the outcome: *Y* = 0.1*X*^2^ + *U* + *ϵ*_*Y*_.
3. Threshold causal effect of the exposure on the outcome: 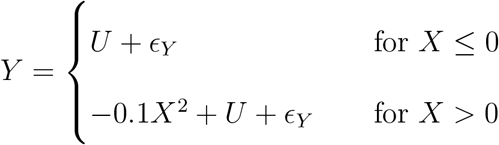

In the first case, the causal effect of the exposure on the outcome is zero throughout. In the second case, the causal effect is negative for negative exposure values and positive for positive exposure values. In the third case, the causal effect is zero for negative exposure values and negative for positive exposure values.

We also consider four models for the effect of the instrument on the exposure:

A. Linearity and homogeneity: *X* = 0.5*Z* + *U* + *ϵ*_*X*_.
B. Non-linearity and homogeneity: 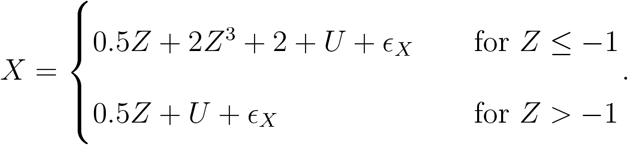
C. Linearity and heterogeneity: *X* = −10 + (1.5 + 0.4*U*)*Z* + *U* + *ϵ*_*X*_
D. As scenario A, but the exposure is coarsened by being rounded to the nearest integer value.

In model A, the effect of the instrument is linear and homogeneous. In model B, it is non-linear as there is a power change at *Z* = −1. In model C, it is heterogeneous, with the effect depending on the value of the confounder *U*. In model D, the effect is linear and homogeneous for the original values of the exposure, but non-homogeneous for the coarsened values.

We consider 12 scenarios, comprising all combinations of exposure–outcome relationships and instrument–exposure models. We simulate data on 10 000 individuals in each dataset, and consider 1000 simulated datasets per scenario. In each scenario, we simulate variables from independent normal distributions: *Z* ∼ 𝒩(0, 0.5^2^), *U* ∼ 𝒩(0, 1^2^), *ϵ*_*X*_ ∼ 𝒩(0, 1^2^), and *ϵ*_*Y*_ ∼ 𝒩(0, 1^2^), where *ϵ*_*X*_ and *ϵ*_*Y*_ are independent error terms for the exposure *X* and outcome *Y* respectively. The average proportion of variation in the exposure explained by the instrument in each scenario is roughly 5%. We divide each dataset into 10 strata of 1000 individuals using both of the stratification methods, and calculate LACE estimates in each stratum. We also assess genetic associations with the exposure in each stratum to see whether these follow the expected pattern.

### Applied investigation: alcohol intake and systolic blood pressure

We implement the residual and doubly-ranked stratification methods to study the causal effect of alcohol intake on SBP. Previous Mendelian randomization analyses have suggested that alcohol intake has positive causal effects on hypertension risk and systolic blood pressure^23–25^, although the shape of the causal relationship has not been considered using this approach. We take data from UK Biobank, a prospective cohort study of around half a million UK residents aged 40 to 69 years at baseline, recruited in 2006-2010 from across the United Kingdom^26^. We consider 385 000 unrelated individuals of European ancestries, who passed various quality control filters as described previously^27^. A small number of individuals (2067) were dropped from the analysis at random to obtain equally-sized strata, facilitating comparison of stratum-specific estimates for this illustrative example.

We first construct a weighted genetic score for each individual from 93 genetic variants which have previously been shown to be associated with alcohol intake in 941 280 individuals at a genomewide level of statistical significance^28^. Only around 124 590 individuals in this analysis were UK Biobank participants, minimizing bias due to sample overlap^29^. The weighted genetic score was centred to have mean zero, so that the mean of the residual exposure was the same as the mean of the exposure. Alcohol assumption was calculated for each participant based on self-reported data on consumption frequency for various alcoholic drinks as described previously^30^, and is measured in units of g/day. We obtain LACE estimates using each stratification method for 77 strata, each of which contains 5000 individuals. These are plotted against the average value of the exposure in each stratum, thus providing insight into the shape of the causal relationship. We also provide LACE estimates for tenths of the population.

## Results

### Simulation study

Results from the residual stratification and the doubly-ranked method are displayed in Figures 3-6. For each scenario, we provide a boxplot of LACE estimates in each stratum. Median estimates from the residual stratification method are close to the average true effect values for that stratum in Scenarios A1, A2, and A3, where the effect of the instrument on the exposure is linear and homogeneous. However, in other scenarios, there is notable bias and distortion of the shape of the exposure-outcome causal relationship, even in Scenarios B1 and C1, where the true causal effect is null, and in Scenario D1, where the effect of the instrument on the exposure is linear and homogeneous, but the exposure is rounded to the nearest integer. The imprecise estimates in Scenario D1 correspond to strata where the majority of people have similar estimates for the exposure, and so the genetic association with the exposure in that stratum is weak. In contrast, median estimates from the doubly-ranked stratification method are close to the true values throughout.

**Figure 3:**
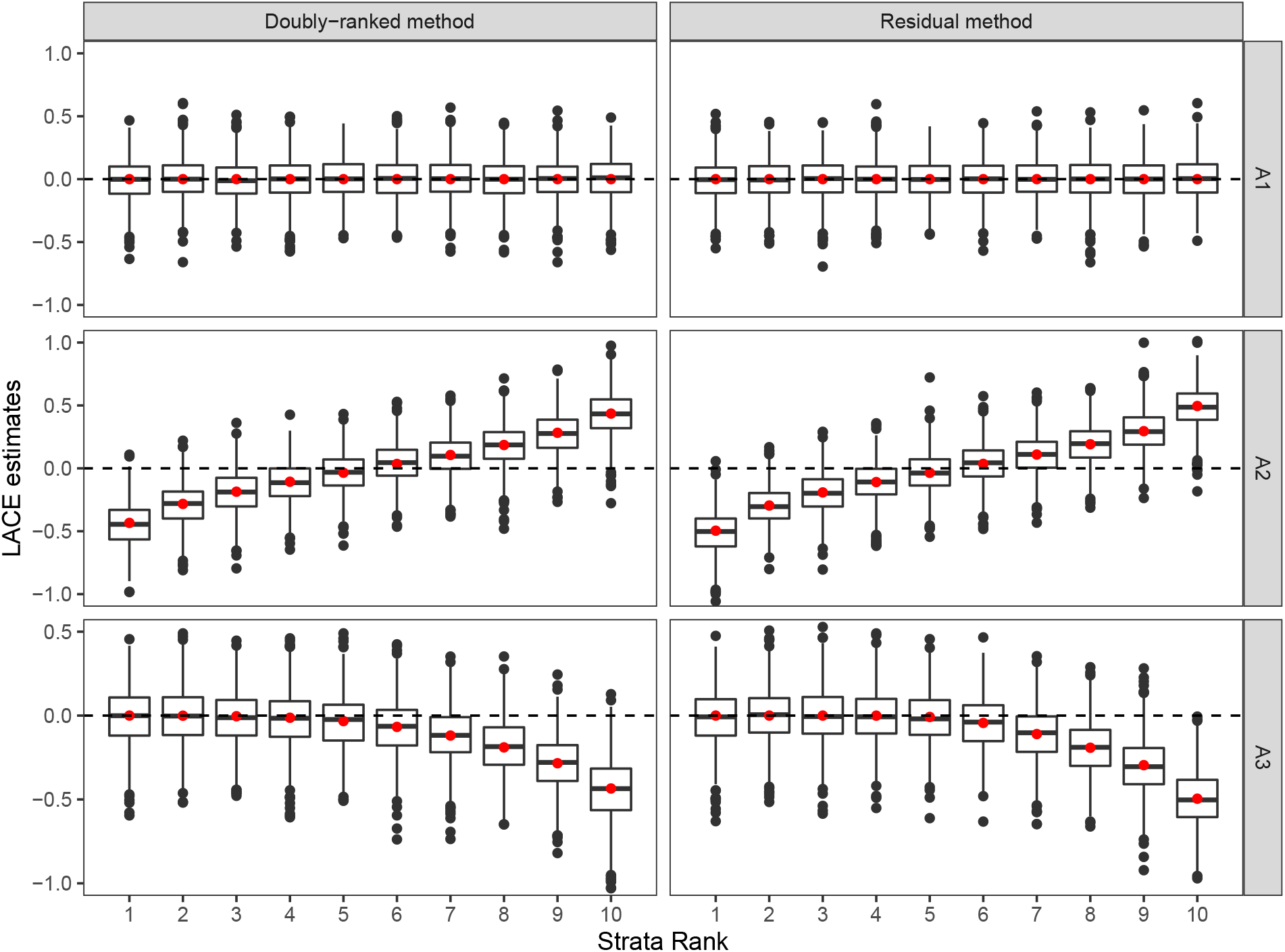
Results of the doubly-ranked method and residual method for model A (linearity and homogeneity) with three different causal relationship between the exposure and the outcome (denoted by A1, A2, A3). Boxplot results represent the LACE estimates within the 10 strata. Red points represent the target causal effects within strata. Box indicates lower quartile, median, and upper quartile; error bars represent the minimal and maximal data point falling in the 1.5 interquartile range distance from the lower/upper quartile; estimates outside this range are plotted separately.

**Figure 4:**
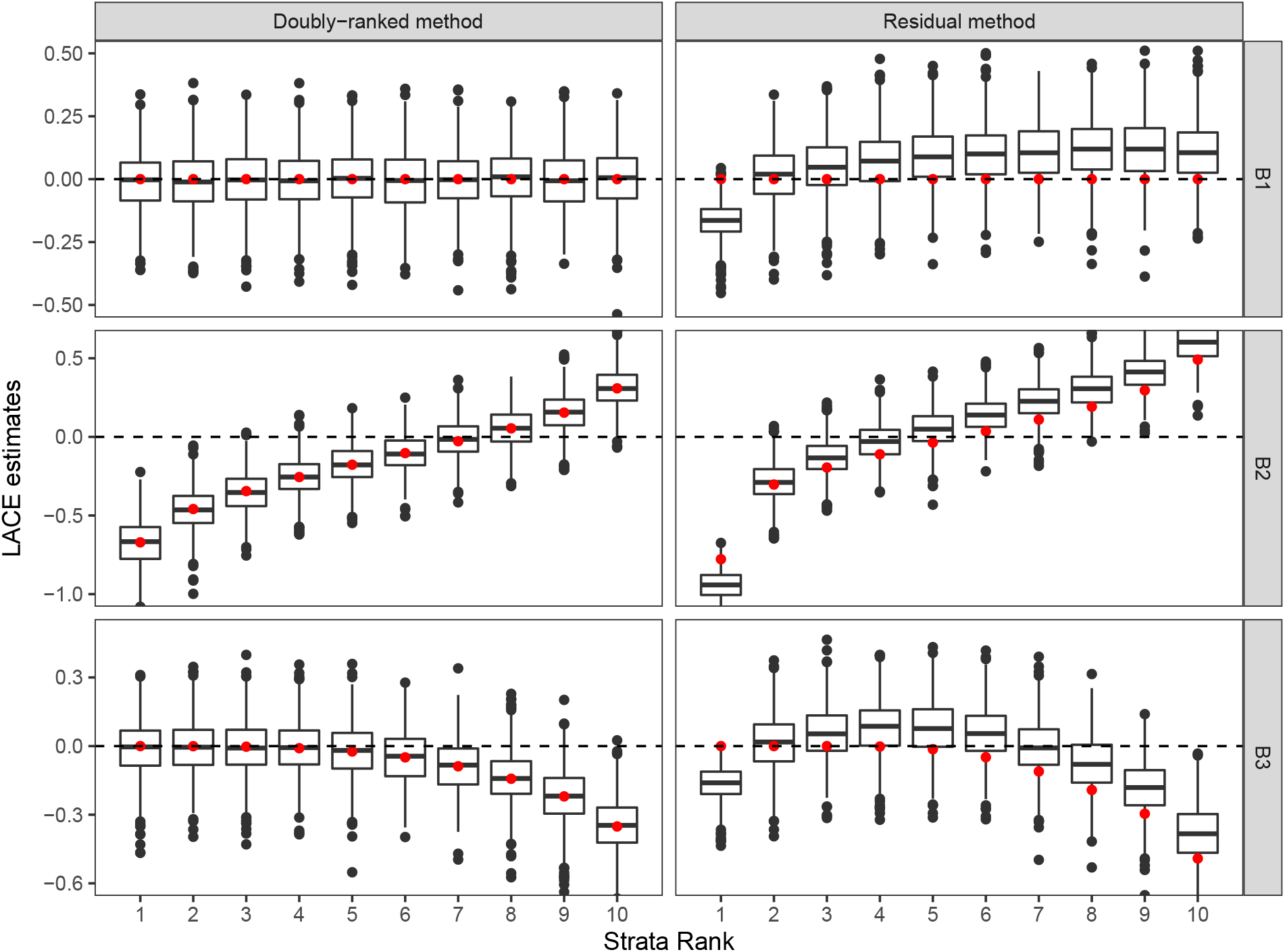
Results of the doubly-ranked method and residual method for model B (nonlinearity and homogeneity) with three different causal relationship between the exposure and the outcome (denoted by B1, B2, B3). Boxplot results represent the LACE estimates within the 10 strata. Red points represent the target causal effects within strata. Box indicates lower quartile, median, and upper quartile; error bars represent the minimal and maximal data point falling in the 1.5 interquartile range distance from the lower/upper quartile; estimates outside this range are plotted separately.

**Figure 5:**
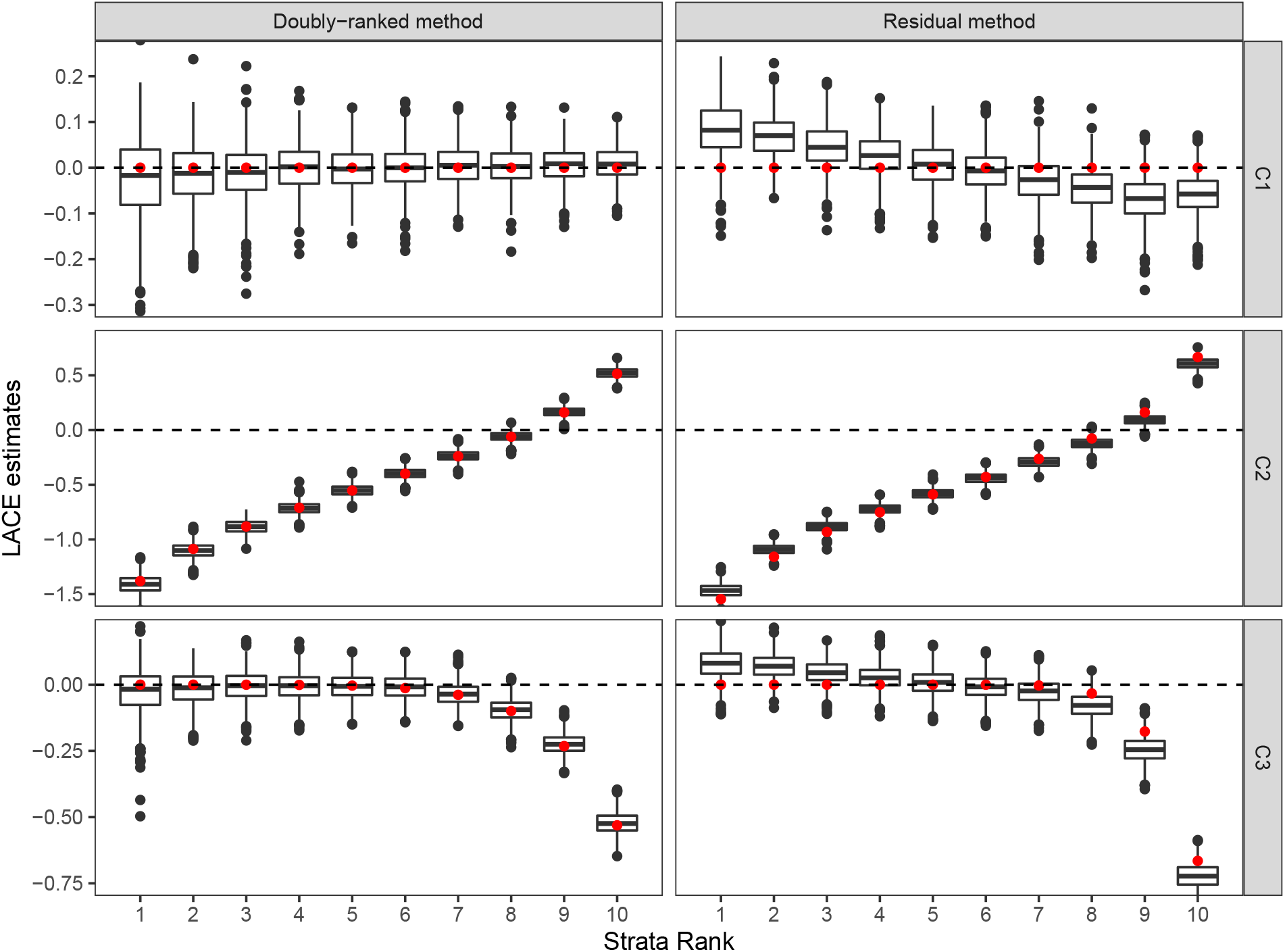
Results of the doubly-ranked method and residual method for model C (linearity and heterogeneity) with three different causal relationship between the exposure and the outcome (denoted by C1, C2, C3). Boxplot results represent the LACE estimates within the 10 strata. Red points represent the target causal effects within strata. Box indicates lower quartile, median, and upper quartile; error bars represent the minimal and maximal data point falling in the 1.5 interquartile range distance from the lower/upper quartile; estimates outside this range are plotted separately.

**Figure 6:**
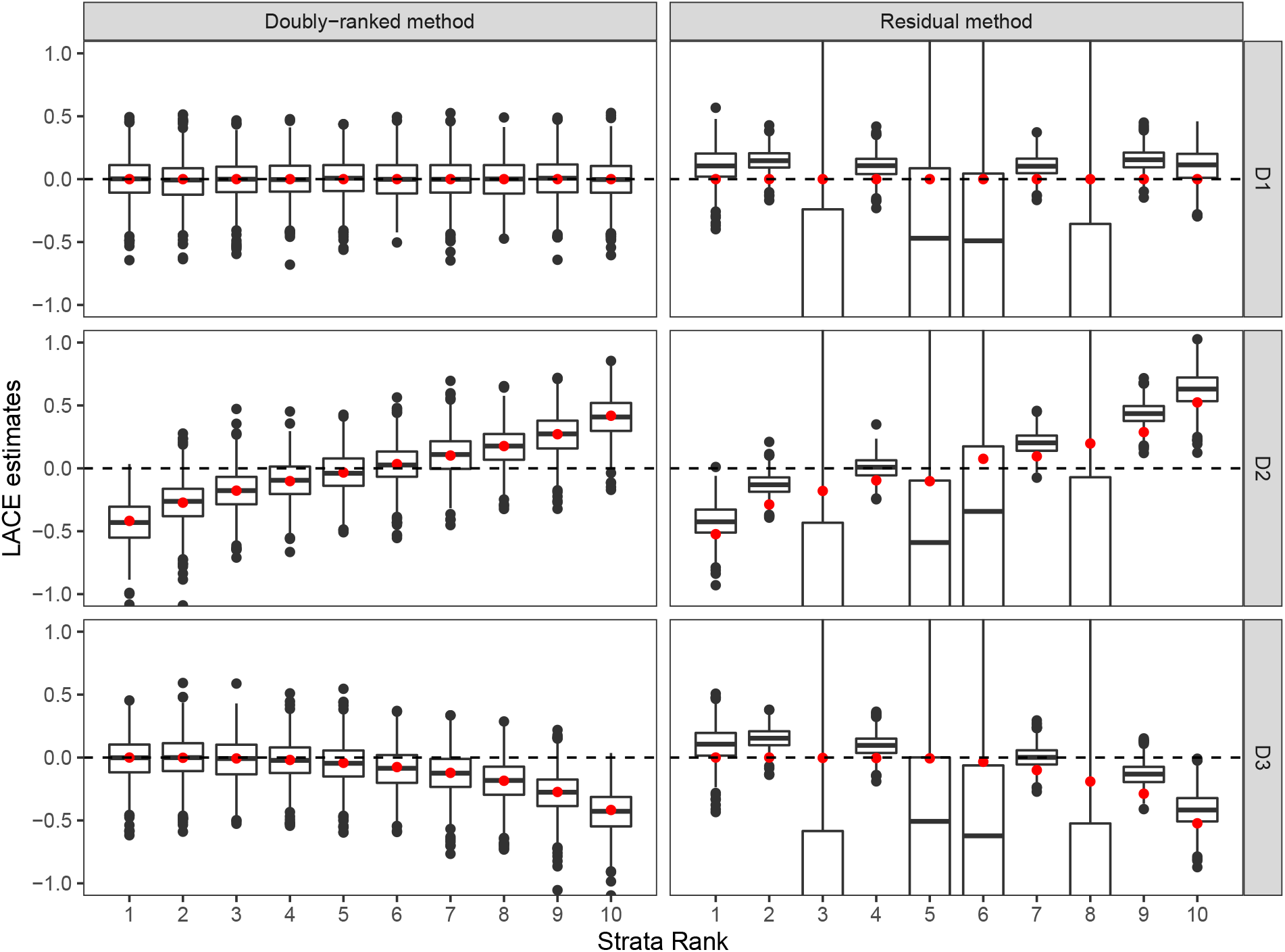
Results of the doubly-ranked method and residual method for model D (coarsened exposures) with three different causal relationship between the exposure and the outcome (denoted by D1, D2, D3). Boxplot results represent the LACE estimates within the 10 strata. Red points represent the target causal effects within strata. Box indicates lower quartile, median, and upper quartile; error bars represent the minimal and maximal data point falling in the 1.5 interquartile range distance from the lower/upper quartile; estimates outside this range are plotted separately.

Table 1 provides two further summaries of the simulation results: the mean squared error (MSE) of estimates summed across the 10 strata, and the coverage of estimates, representing the proportion of LACE estimates for which the 95% confidence interval included the average causal effect for that stratum, defined as the weighted integral of the derivative of the function relating the exposure to the outcome with the weight function estimated by the instrument and exposure values of individuals in each stratum (as shown and derived in Text S2). We refer to this average causal effect as the target causal effect. When the effect of the instrument on the exposure is linear and homogeneous (model A), both methods perform similarly. For other models, the doubly-ranked method has better performance than the residual method in terms of both MSE and coverage. For models B and C, the residual method has inflated type I error rates while the doubly-ranked method has appropriate coverage rates. In addition, the doubly-ranked method has prominently better performance in the coarsened exposure case (model D), whereas residual methods do not work well even under the linearity and homogeneity model. We note that the Gelman–Rubin uniformity statistic was below the threshold value of 1.02 in all strata for the coarsened exposure (see Text S1).

**Table 1:** Summary of simulation study results: mean squared errors (MSE) and coverage of the 95% confidence interval for the residual and doubly-ranked stratification method in each scenario. MSE is summed across estimates in all 10 strata, and results are averaged across 1000 datasets per scenario.

| Scenario | Residual method |  | Doubly-ranked method |  |
| --- | --- | --- | --- | --- |
|  | MSE | Coverage | MSE | Coverage |
| A1 | 0.025 | 0.947 | 0.026 | 0.957 |
| A2 | 0.025 | 0.949 | 0.026 | 0.957 |
| A3 | 0.026 | 0.947 | 0.027 | 0.952 |
| B1 | 0.023 | 0.810 | 0.014 | 0.949 |
| B2 | 0.023 | 0.818 | 0.013 | 0.958 |
| B3 | 0.024 | 0.808 | 0.014 | 0.951 |
| C1 | 0.005 | 0.805 | 0.003 | 0.950 |
| C2 | 0.005 | 0.811 | 0.003 | 0.959 |
| C3 | 0.005 | 0.803 | 0.003 | 0.953 |
| D1 | 2.681 | 0.785 | 0.027 | 0.953 |
| D2 | 2.941 | 0.794 | 0.027 | 0.955 |
| D3 | 3.500 | 0.790 | 0.028 | 0.951 |

Additionally, we assessed the genetic associations with the exposure within strata defined by the two methods. Results are shown in Supplementary Figures S1 to S4. Genetic associations with the exposure in strata defined by the doubly-ranked method follow the expected pattern: homogeneous for models A, B, and D, but monotone increasing for model C. In contrast, genetic associations with the exposure in strata defined by the residual method are similar for strata 2 to 9 in all non-coarsened exposure scenarios, even for model C, where they should not be similar. With a coarsened exposure (model D), genetic associations with the exposure in strata defined by the residual method are highly irregular, providing further evidence on the unreliability of this method in the coarsened exposure scenario. Our results suggest the approach of assessing stratum-specific genetic associations with the exposure is only reliable as a test of homogeneity of the instrument effect using the doubly-ranked method.

To assess the performance of the method with a more limited instrument, we repeated the simulation study under model A except that the genetic instrument only took two values: *Z* = 0, 1. This was achieved by simulating from a Bernoulli distribution with probability 0.3. Results are shown in Supplementary Figure S5. We can see that the doubly-ranked method worked equally well in this scenario.

### Applied investigation: alcohol intake and systolic blood pressure

We proceed to consider the effect of alcohol intake on SBP using data from UK Biobank. Even though the level of alcohol intake is a continuous measure, it has a natural left truncation at zero g/day. Additionally, while detailed information on various categories of alcohol consumption is available in UK Biobank, the questionnaire nature of the data means that several individuals have the same reported value for alcohol intake. Overall, 5284 unique values of alcohol intake were represented in the data, and 21 340 individuals reported zero alcohol intake. This therefore represents a coarsened exposure scenario. LACE estimates represent change in SBP in mmHg units per 1 g/day increase in genetically-predicted values of alcohol intake.

Graphs of LACE estimates from the two stratification methods plotted against average alcohol intake in that stratum are presented in Figure 7. For the residual stratification method, most estimates in the low alcohol range are in the negative direction. But residual alcohol is negative for some individuals with zero alcohol consumption; this value does not have a natural interpretation. It is logically impossible that the genetic effect on alcohol consumption is homogeneous in the population. The effect of the genetic variants for zero consumption individuals cannot be positive, as this would imply negative consumption if they had a different value of the genetic variants. Therefore, estimates in low consumption strata from the residual method are unreliable.

**Figure 7:**
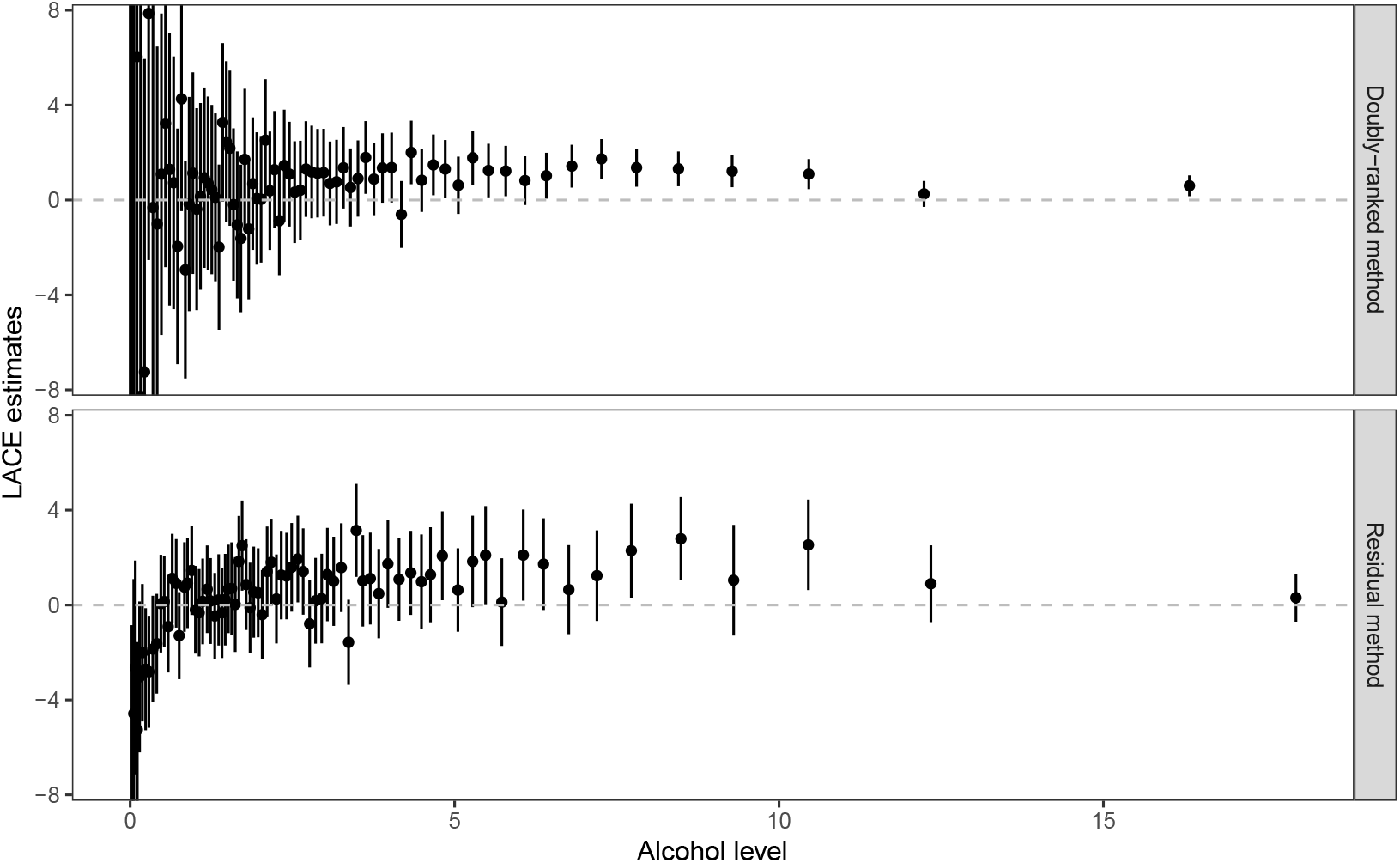
LACE estimates of alcohol intake on SBP from the two stratification methods (residual method and doubly-ranked method) against average levels of alcohol intake in the 77 strata. The error bars represent the 95% confidence interval for each stratum-specific estimate.

**Figure 8:**
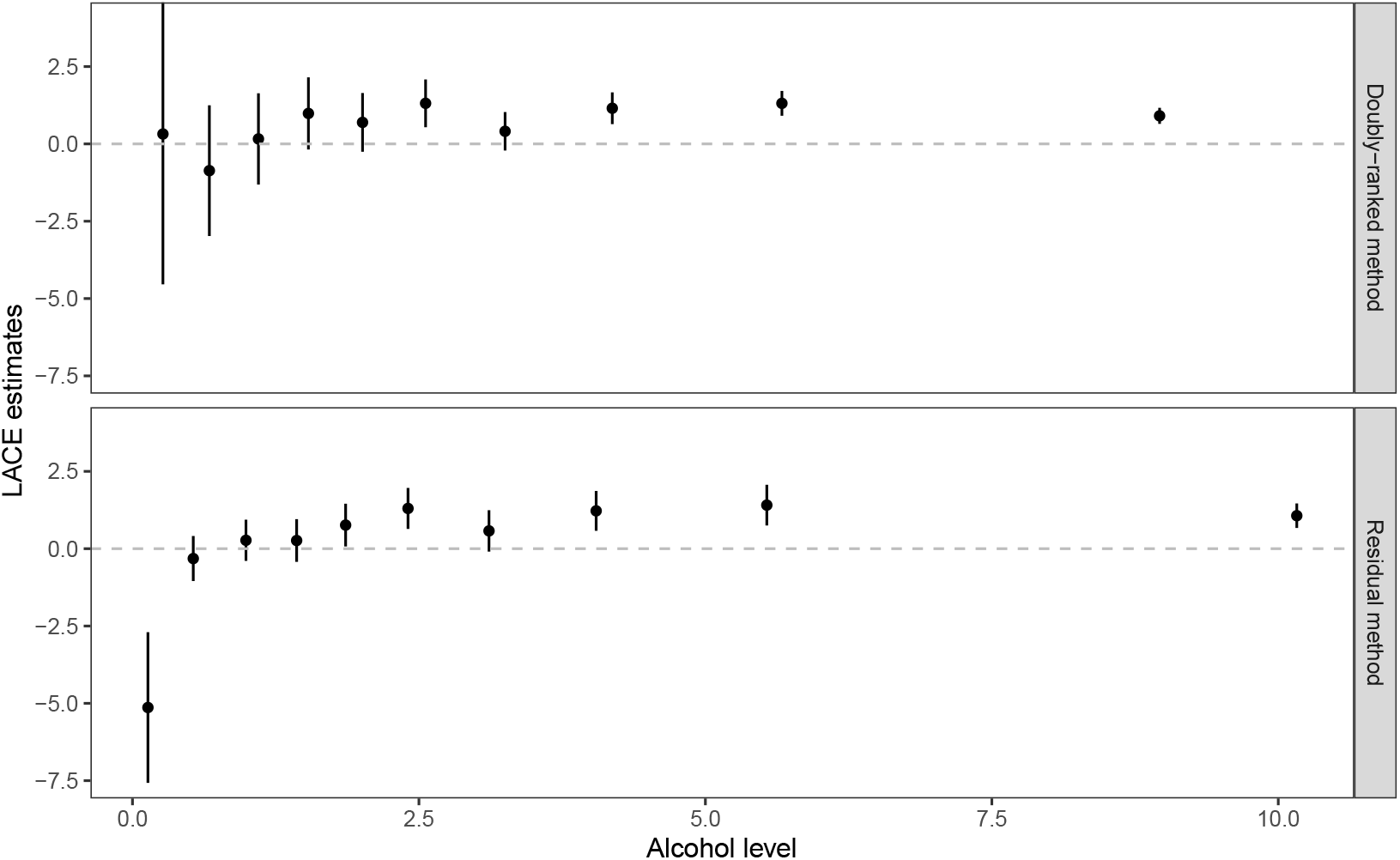
LACE estimates of alcohol intake on SBP from the two stratification methods (residual method and doubly-ranked method) against average levels of alcohol intake in the 10 strata. The error bars represent the 95% confidence interval for each stratum-specific estimate.

In contrast, LACE estimates from the doubly-ranked method are generally positive throughout. For moderate to heavy drinkers (*>*5 g/day alcohol intake), estimates from the two methods are similar, but those from the doubly-ranked method have less uncertainty and a greater proportion of 95% confidence intervals that exclude the null. This may be due to the increasing genetic associations with the exposure within strata for the doubly-ranked method, leading to more precise ratio estimates as observed for model C of the simulation study.

A similar pattern of LACE estimates is seen when dividing the population into tenths using both methods: the LACE estimate is −5.13 (95% confidence interval −7.57 to −2.70) in the lowest decile group for the residual method, and 0.32 (95% confidence interval −4.54 to 5.18) for the doubly ranked method (Table 2). The negative estimate in the lowest quantile for the residual method is biologically implausible. It is also implausible that the causal estimate is stronger in the lowest decile group than in the other groups. In contrast, the estimates from the doubly-stratified method are close to the null in the lowest quantiles, and suggest a positive effect from the fourth quantile onwards, with significant positive estimates in strata 6, 8, 9, and 10 (*p <* 0.05). This is biologically plausible, as alcohol will remain in the system longer for moderate to heavy drinkers, and so the effect of a 1 g/day increase will likely be greater than for light drinkers^31^. Moderate to heavy drinkers are also more likely to include episodic drinkers (so called ‘binge drinkers’), for whom alcohol potentially has a more strongly harmful effect^32^.

**Table 2:** Summary of applied investigation results: the mean alcohol intake (g/day) with its interquartile interval, LACE estimate and 95% confidence interval (mmHg change in systolic blood pressure per 1 g/day increase in genetically-predicted alcohol intake) in each stratum using the residual method and doubly-ranked method. IQR represents the interval for the interquartile range in each stratum. CI represents the 95% confidence interval.

| Stratum | Residual method |  |  | Doubly-ranked method |  |  |
| --- | --- | --- | --- | --- | --- | --- |
|  | Mean (IQR) | Estimate | 95% CI | Mean (IQR) | Estimate | 95% CI |
| 1 | 0.13(0.00,0.24) | -5.14 | (-7.57,-2.70) | 0.27(0.00,0.45) | 0.32 | (-4.54,5.18) |
| 2 | 0.53(0.33,0.77) | -0.32 | (-1.05,0.41) | 0.67(0.24,1.03) | -0.87 | (-2.98,1.25) |
| 3 | 0.99(0.77,1.21) | 0.27 | (-0.39,0.94) | 1.10(0.65,1.51) | 0.16 | (-1.31,1.64) |
| 4 | 1.43(1.29,1.56) | 0.26 | (-0.43,0.95) | 1.54(1.03,1.93) | 0.98 | (-0.18,2.15) |
| 5 | 1.86(1.69,2.06) | 0.76 | (0.07,1.45) | 2.01(1.47,2.46) | 0.70 | (-0.25,1.64) |
| 6 | 2.40(2.21,2.60) | 1.30 | (0.64,1.96) | 2.56(1.83,3.09) | 1.31 | (0.54,2.08) |
| 7 | 3.11(2.89,3.35) | 0.58 | (-0.09,1.24) | 3.25(2.31,3.94) | 0.41 | (-0.21,1.03) |
| 8 | 4.05(3.71,4.40) | 1.22 | (0.58,1.87) | 4.19(3.09,5.09) | 1.15 | (0.64,1.67) |
| 9 | 5.54(5.09,6.04) | 1.41 | (0.75,2.07) | 5.67(4.00,6.86) | 1.31 | (0.91,1.71) |
| 10 | 10.16(7.60,11.41) | 1.07 | (0.67,1.46) | 8.96(5.86,10.91) | 0.91 | (0.65,1.17) |

Genetic associations with the exposure in strata are displayed in Supplementary Figure S6. While genetic associations in strata defined by the residual method were similar for strata 2 to 9, we showed in the simulation study that this is not a reliable way of assessing variability in instrument strength. In contrast, genetic associations with the exposure in strata defined by the doubly-ranked method were monotone increasing across the strata. This suggests the genetic variants have a stronger effect on alcohol intake for higher levels of alcohol consumption. This provides further empirical evidence that the doubly-ranked method is more appropriate for this example. Values of the Gelman–Rubin uniformity statistic were below 1.02 for both the stratification into 77 strata and into 10 strata, suggesting that the number of strata is not excessive given the degree of coarsening. The distribution of the exposure in strata defined by the doubly-ranked method is normally broader than that by the residual method.

A potential limitation of the analysis is that the zero alcohol consumption group contains both never-drinkers and ex-drinkers. However, in contrast to chronic disease outcomes, the impact of alcohol consumption on SBP is likely to be short-term in nature^33^. Therefore any lifelong effect of exposure to alcohol in ex-drinkers is less likely to impact findings for this outcome.

## Discussion

Non-linear Mendelian randomization is an extension to standard Mendelian randomization that stratifies the population into subgroups with different average levels of the exposure, and then performs separate instrumental variable analyses in each stratum of the population. In this paper, we have proposed the doubly-ranked method, a non-parametric method for constructing strata of the population, such that the strata are uncorrelated with the instrument, and the average level of the exposure is increasing across strata. In the case where the instrument takes a fixed value in each pre-stratum, the correlation between the instrument and stratification is exactly zero. In other cases, this correlation should be close to zero. In either case, stratification using this method should not induce substantial collider bias, and so if the genetic instrument is valid in the population as a whole, it should be valid in strata of the population defined using the doubly-ranked method.

We validated the doubly-ranked method in a simulation study for a range of scenarios, including various models for the effect of the exposure on the outcome (null, U-shaped, threshold) and for the effect of the instrument on the exposure. The previously proposed residual stratification method provided unbiased estimates when the effect of the instrument on the exposure was linear and homogeneous, but otherwise provided biased estimates with inflated Type I error rates. In contrast, the doubly-ranked stratification method provided unbiased estimates and appropriate coverage rates when the effect of the instrument was non-linear or heterogeneous. It also provided unbiased estimates when the exposure was coarsened by rounding its value to the nearest integer; another scenario that can lead to large bias for the residual method. The coarsened exposure scenario is particularly important in applied practice, as many exposures are reported as rounded values or in categories. We also assessed the performance of the doubly-ranked method in an applied example, which demonstrated evidence for an effect of alcohol on systolic blood pressure that was positive across strata at moderate to high levels of alcohol consumption, while the residual method suggested a negative effect at low levels of alcohol consumption that is not biologically plausible.

Aside from relaxing the parametric assumptions required by the residual method, the doubly-ranked method is able to deal with a wider set of exposures. A further feature is that strata formed by the doubly-ranked method are equally-sized, meaning that the precision of LACE estimates should be similar across the distribution of the exposure. A disadvantage of the method is that it is harder to stratify the population according to clinically-defined threshold values of the exposure, such as the World Health Organization categories for BMI (BMI *<* 18.5 kg/m^2^ is underweight, and so on). Similarly, it is difficult to define which individuals are selected into each stratum, and so whom each stratum-specific estimate relates to.

In addition to being used to estimate the non-linear relationship, the doubly-ranked method can also assess the homogeneity assumption. Under homogeneity, the genetic associations with the exposure in strata obtained from the doubly-ranked method should be consistent to a fixed value. Any inconsistent pattern, like the increasing pattern for the alcohol example, indicates the effect of the instrument on the exposure is heterogenous, and hence the doubly-ranked method should be used in preference to the residual method.

Previous work has considered smoothing LACE estimates from the residual method using either a fractional polynomial or piecewise linear method^4^. These methods could equally be applied to LACE estimates from the doubly-ranked method. The result of these methods is a plot of the outcome against the exposure, which can more easily be compared to the shape of the association estimated in a conventional observational epidemiological analysis. The LACE estimates represent the gradient of this curve, as they represent the average causal effect at that level of the exposure, and so the plot of the LACE estimates against the exposure is the derivative of the plot of the outcome against the exposure. A plot of the LACE estimates against the exposure is less conventional, but it more clearly represents the quantities estimated in a non-linear Mendelian randomization investigation, which are the stratum-specific causal effects.

In summary, this paper presents the doubly-ranked stratification method, a method for relaxing the parametric assumptions made in non-linear Mendelian randomization investigations. We recommend analysts consider using this method in preference to the residual stratification method, or at least as a sensitivity analysis to assess the dependence of findings on the linearity and homogeneity assumptions for the effect of the instrument on the exposure.

## Funding

AMM is funded by the EU/EFPIA Innovative Medicines Initiative Joint Undertaking BigData@Heart grant 116074. SB is supported by a Sir Henry Dale Fellowship jointly funded by the Wellcome Trust and the Royal Society (204623/Z/16/Z). This research was supported by core funding from the: United Kingdom Research and Innovation Medical Research Council (MC_UU_00002/7), British Heart Foundation (RG/13/13/30194; RG/18/13/33946) and NIHR Cambridge Biomedical Research Centre (BRC-1215-20014) [*]. *The views expressed are those of the author(s) and not necessarily those of the NIHR or the Department of Health and Social Care.

### Acknowledgements

The UK Biobank study has approval from the North West Multicentre Research Ethics Committee (11/NW/0382). The research has been conducted using the UK Biobank Resource under Application Number 7439. For the purpose of open access, the author has applied a Creative Commons Attribution (CC BY) licence to any Author Accepted Manuscript version arising from this submission.

## Supporting information

The following supporting information is available as part of the online article:

**Figure S1-S4**. Genetic associations with the exposure for each scenario.

**Figure S5**. Results from the residual and doubly-ranked method with binary instrument values.

**Figure S6**. Genetic associations with the exposure for the real example.

**Text S1**. Exchangeability assessment for the coarsened exposure in each stratum.

**Text S2**. Mathematical details for the causal effect in each stratum.

## Supplementary Figure

**Supplementary Figure S1:**
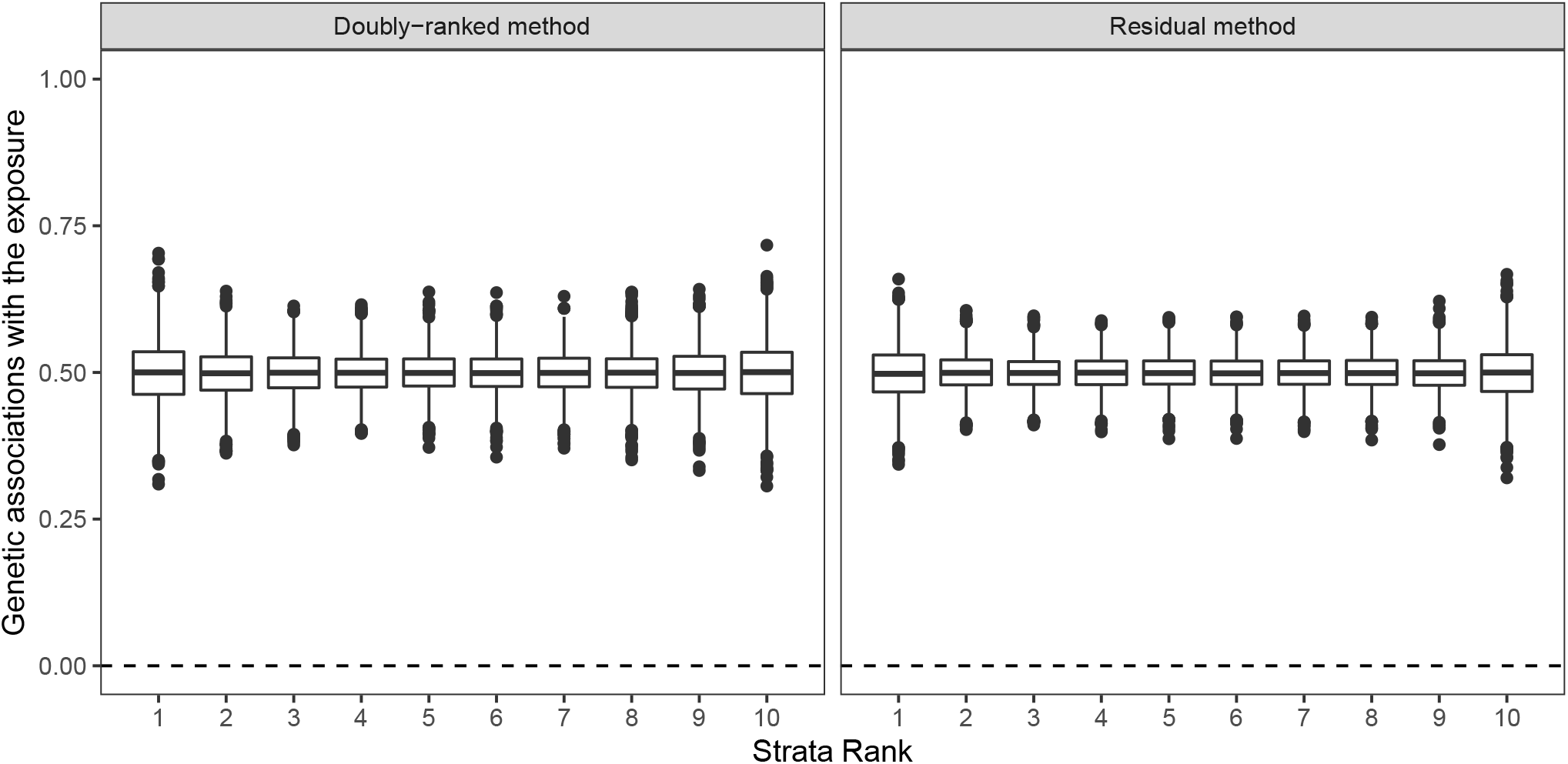
Results of the doubly-ranked method and residual method for model A (linearity and homogeneity). Boxplot results represent the estimates of genetic associations with the exposure within the 10 strata under 3000 simulations. Box indicates lower quartile, median, and upper quartile; error bars represent the minimal and maximal data point falling in the 1.5 interquartile range distance from the lower/upper quartile; estimates outside this range are plotted separately.

**Supplementary Figure S2:**
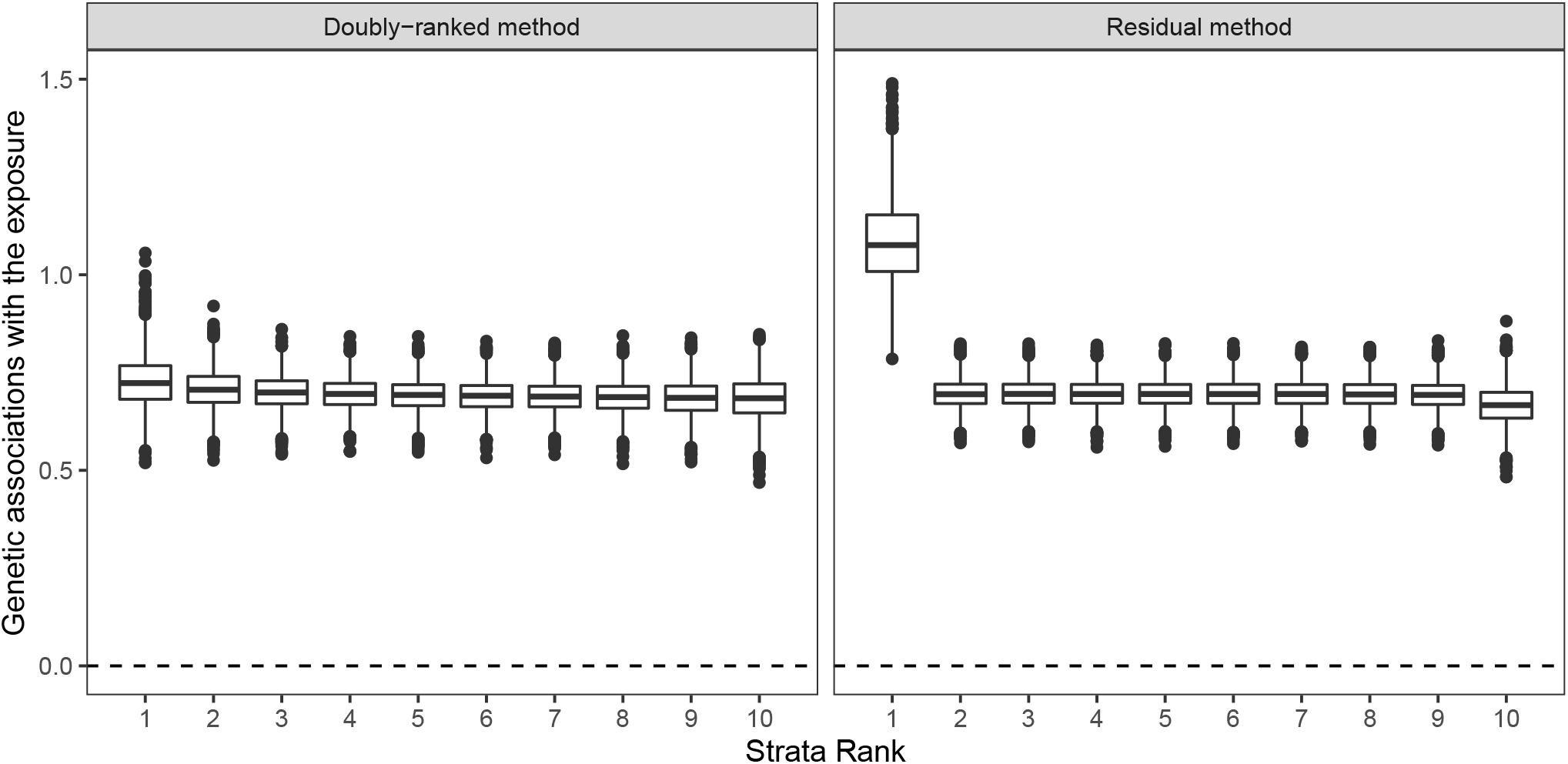
Results of the doubly-ranked method and residual method for model B (nonlinearity and homogeneity). Boxplot results represent the estimates of genetic associations with the exposure within the 10 strata under 3000 simulations. Box indicates lower quartile, median, and upper quartile; error bars represent the minimal and maximal data point falling in the 1.5 interquartile range distance from the lower/upper quartile; estimates outside this range are plotted separately.

**Supplementary Figure S3:**
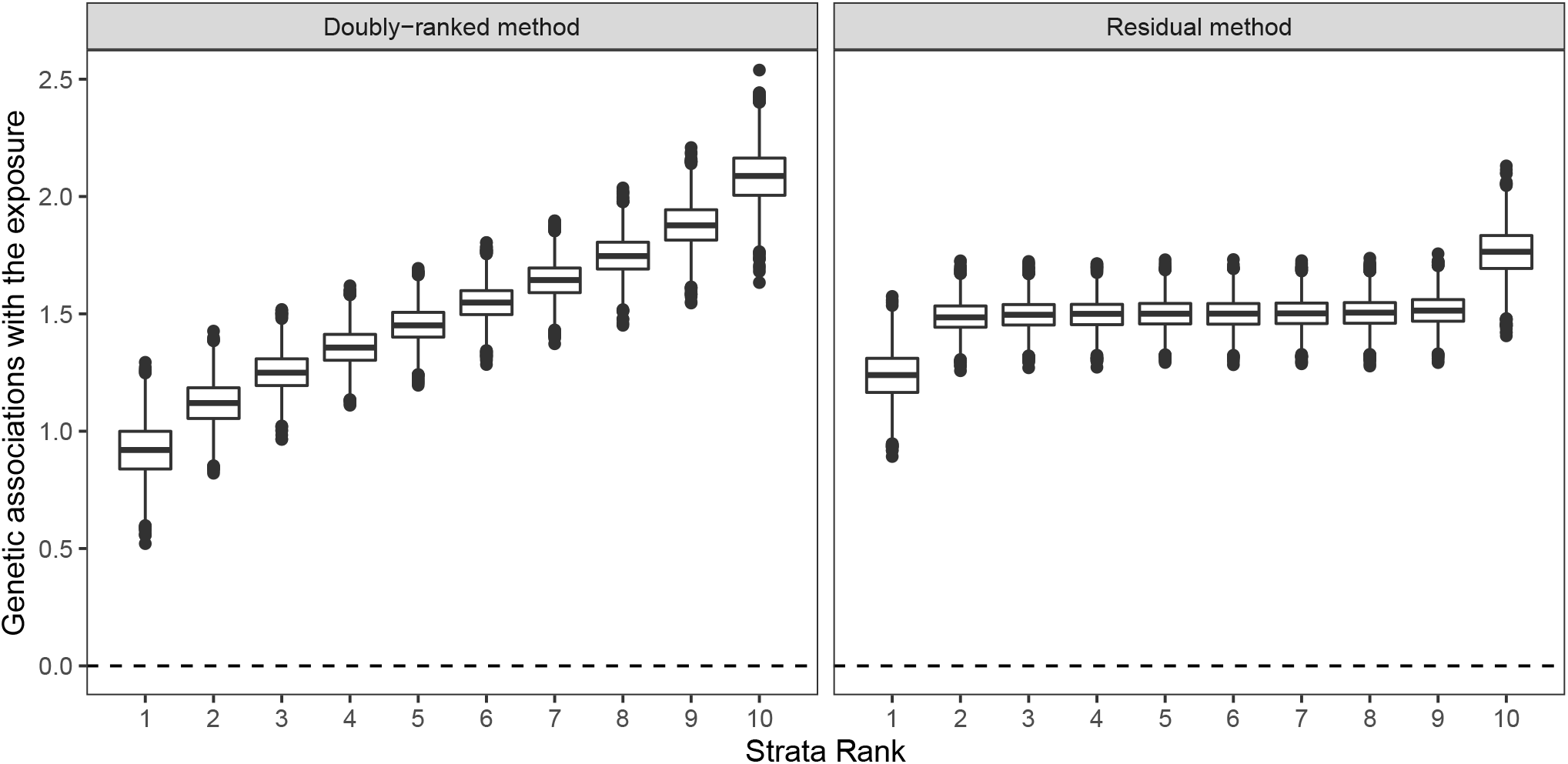
Results of the doubly-ranked method and residual method for model C (linearity and heterogeneity). Boxplot results represent the estimates of genetic associations with the exposure within the 10 strata under 3000 simulations. Box indicates lower quartile, median, and upper quartile; error bars represent the minimal and maximal data point falling in the 1.5 interquartile range distance from the lower/upper quartile; estimates outside this range are plotted separately.

**Supplementary Figure S4:**
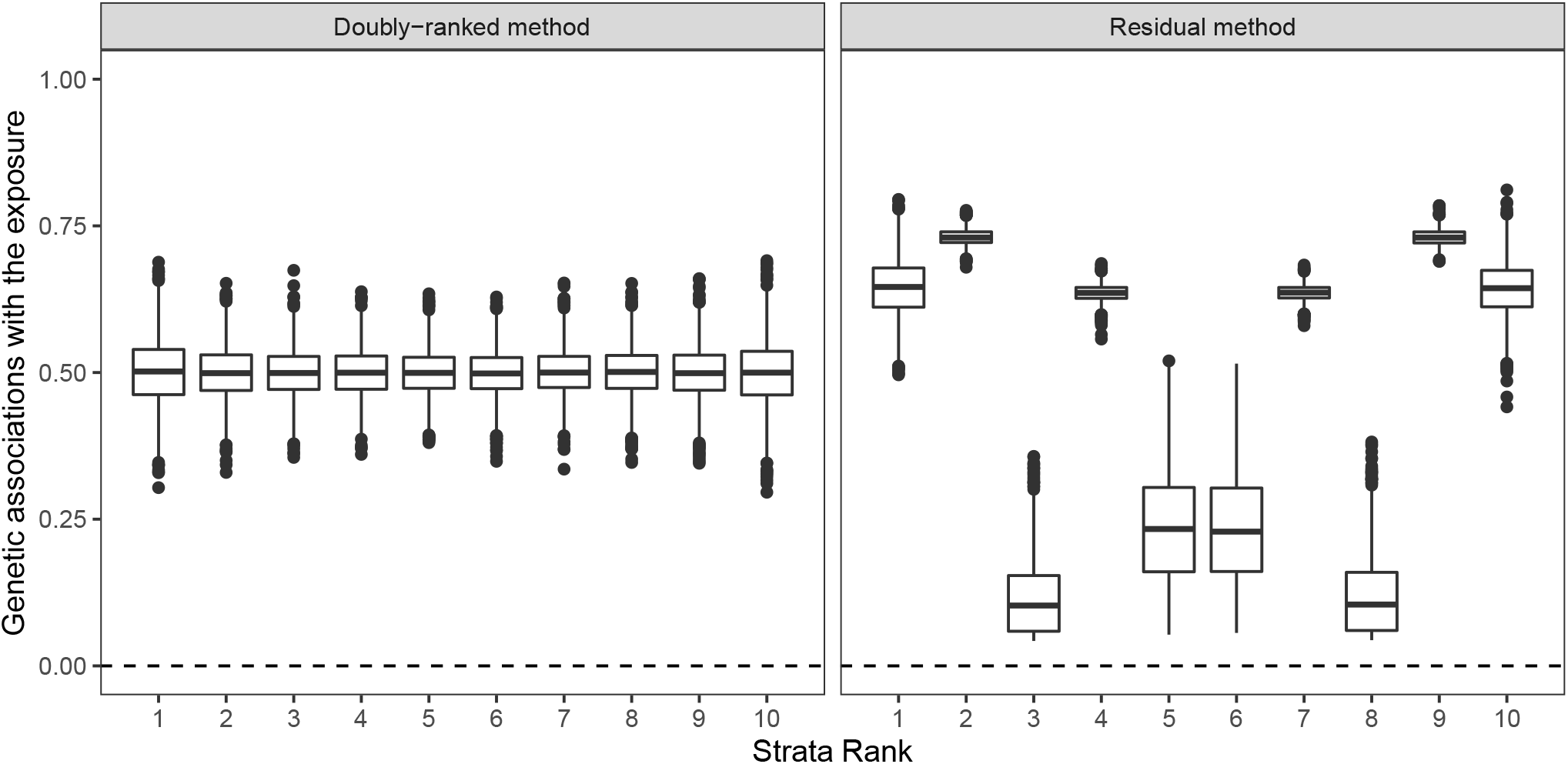
Results of the doubly-ranked method and residual method for model D (coarsened exposures). Boxplot results represent the estimates of genetic associations with the exposure within the 10 strata under 3000 simulations. Box indicates lower quartile, median, and upper quartile; error bars represent the minimal and maximal data point falling in the 1.5 interquartile range distance from the lower/upper quartile; estimates outside this range are plotted separately.

**Supplementary Figure S5:**
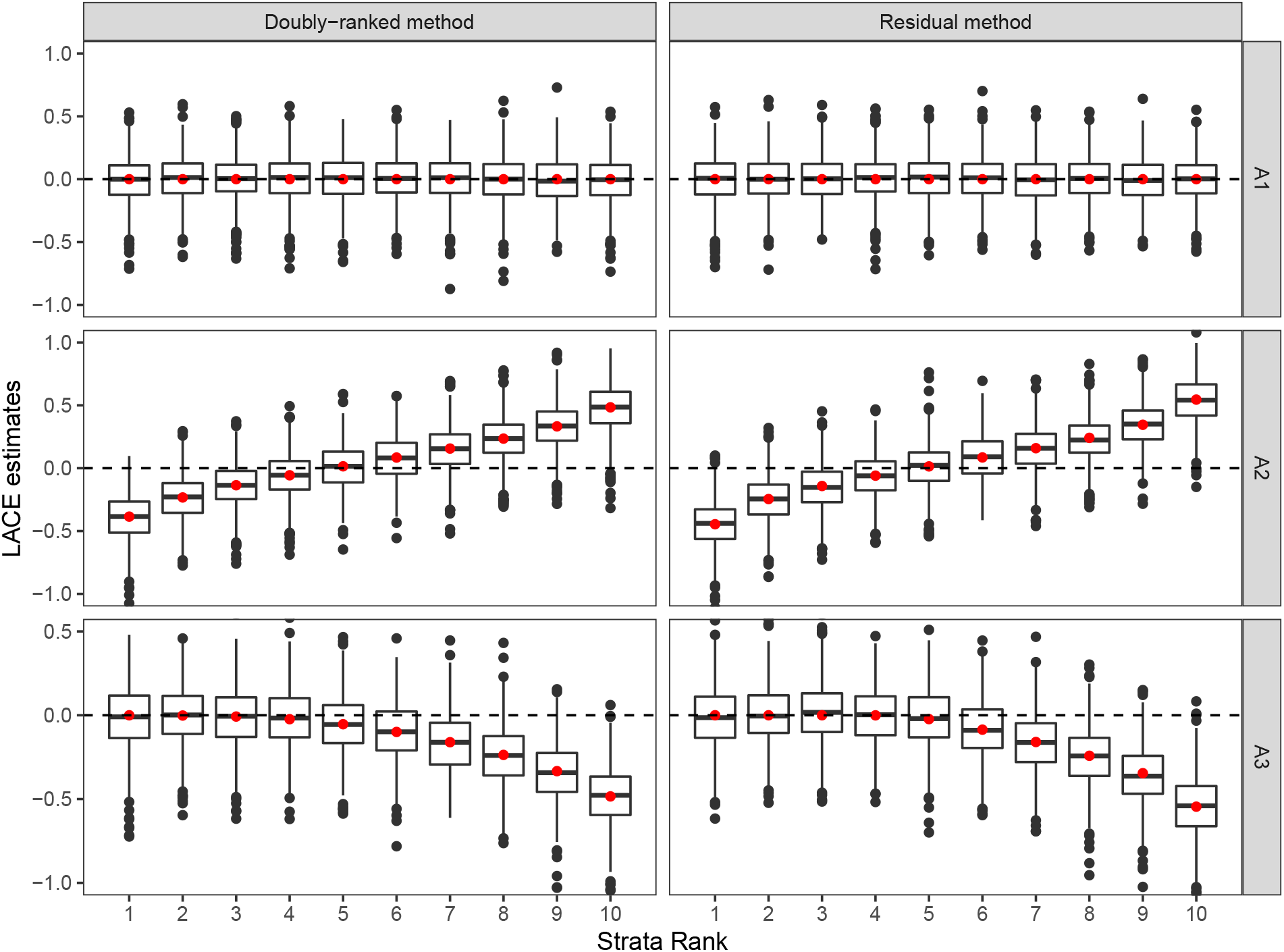
Results of the doubly-ranked method and residual method for model A (homogeneity) with three different causal relationship between the exposure and the outcome (denoted by A1, A2, A3). The instrument follows a Bernoulli distribution with probability 0.3. Boxplot results represent the LACE estimates within the 10 strata. Red points represent the target causal effects within strata. Box indicates lower quartile, median, and upper quartile; error bars represent the minimal and maximal data point falling in the 1.5 interquartile range distance from the lower/upper quartile; estimates outside this range are plotted separately.

**Supplementary Figure S6:**
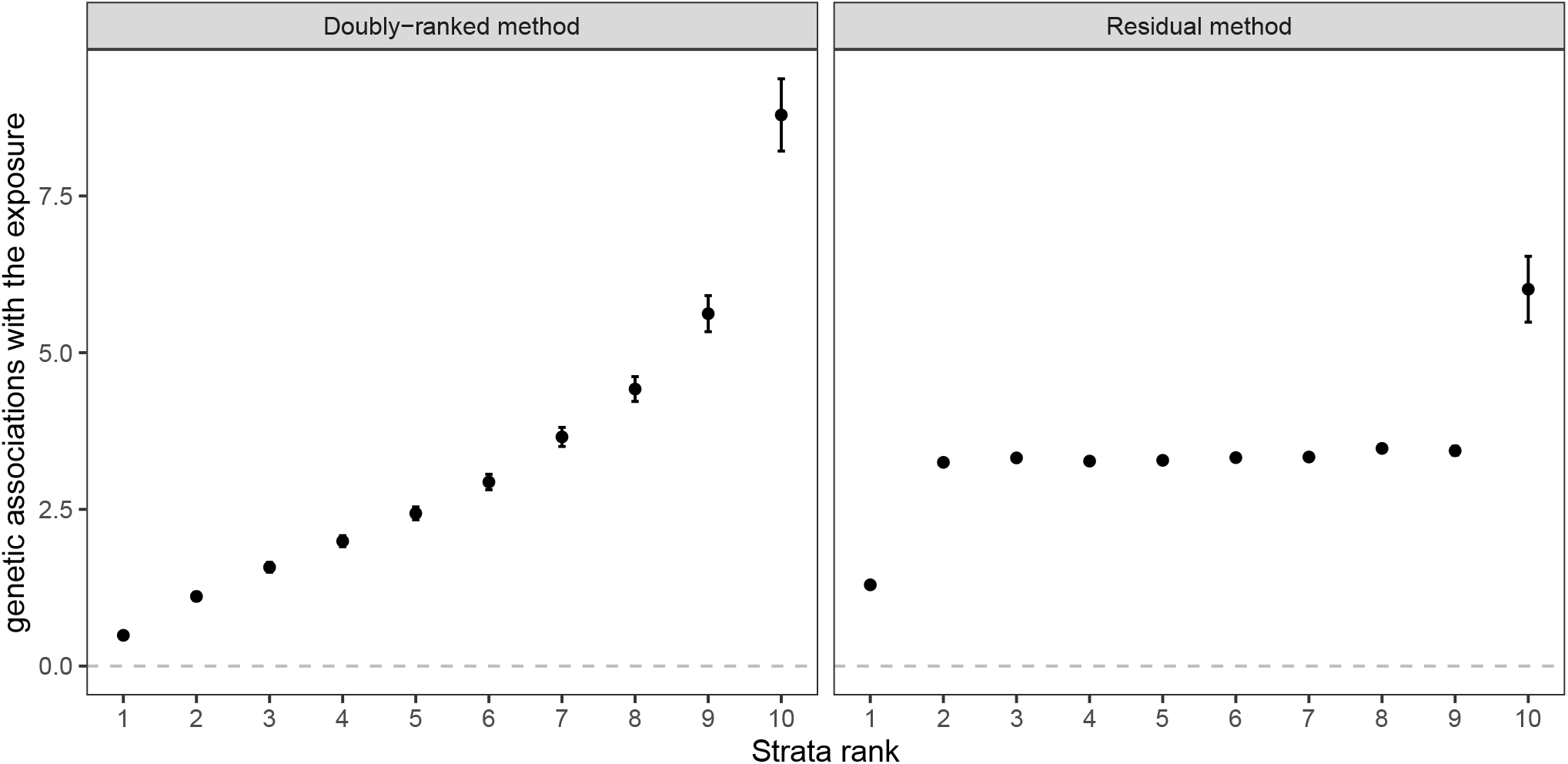
The estimated genetic association with the exposure at each stratum for the residual and doubly-ranked stratification method with the real data of alcohol.

## Text S1: Exchangeability assessment for the coarsened exposure

We give guidance to choose the appropriate number of strata for coarsened exposures in the doubly-ranked stratification method. Let *Z, X* and *U* be the instrument, the (original) exposure, and error term (which incorporates the unmeasured confounding) respectively. Let the observed (coarsened) exposure be *X*^*^ and it satisfies the rank perservation condition that *X*_*i*_ *> X*_*l*_ for any 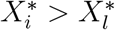. For example, when exposure is coarsened by being rounded to the nearest integer value, *X*_*i*_ *> X*_*l*_ for any [*X*_*i*_] *>* [*X*_*l*_]. Now for simplification assume the instrument *Z* takes *K* different values *J* times. We can build *J* strata via the doubly-ranked method according to the observed exposure *X*^*^. Namely, the *j*-th strata (here we ignore the outcome information) is

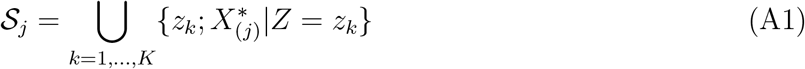

where 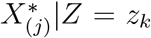 represents the *j*-th ranked exposure at the pre-strata of *Z* = *z*_*k*_. When some observed exposures have the same value as a clump, we will randomly sort them. Due to the rank preservation property, the distribution of the error term at the *k*-th pre-strata (i.e. *Z* = *z*_*k*_) and the *j*-th strata for the coarsened value of the exposure, denoted by 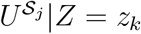, can be expressed as a categorical random variable with the probability mass function

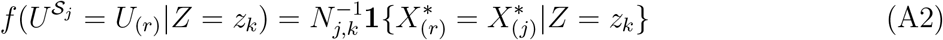

where *N*_*j,k*_ represents the size of the exposure clump in which the *j*-th ranked observed exposure of the *k*-th pre-strata is located. That is,

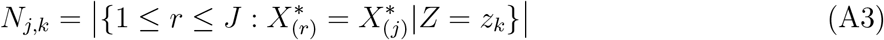

It is easy to know that the distribution of 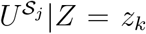 is determined by the two coefficients *a*_*j,k*_ and *b*_*j,k*_

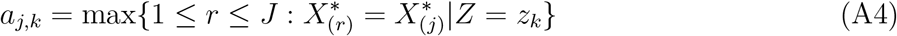

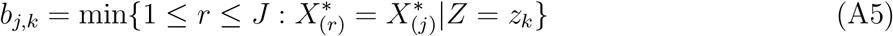

For the *j*-th strata 𝒮_*j*_, the confounding should be approximately independent with the instrument (i.e., exchangeability) when the distribution of 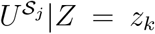 is uncorrelated with *z*_*k*_. That is, {*a*_*j,k*_, *b*_*j,k k*=1,2,…,*K*_ has stabilized distribution over *k*. When exposure is precisely measured (i.e., the original exposure *X*), it is clear that *a*_*j,k*_ = *b*_*j,k*_ = *j* and the confounding is exactly independent of the instrument. The stabilization can be checked in many ways. One heuristic method is the Gelman–Rubin convergence diagnostic, which evaluates whether multiple sample series (e.g. MCMC chain samples) have the converged distribution. In our analysis, we split {*a*_*j,k*}*k*=1,2,…,*K*_ (similar to {*b*_*j,k*}*k*=1,2,…,*K*_) into *N* ^*^ equal parts, each of which is regarded as a chain with the length of *n*^*^ = *K/N* ^*^. We denote the elements of the *t*-th sample at the i-th chain by *x*_*i,t*_, where *i ∈* {1, 2, …, *N* ^*^} and *t ∈* {1, 2, …, *n*^*^}. That is, *x*_*i,t*_ = *a*_*j,n* *(*i* − 1)+*t*_. The between-chain variance estimate is

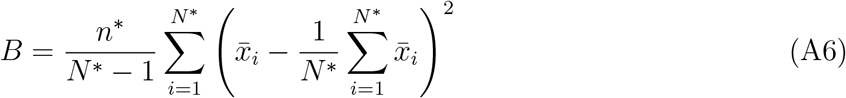

where the mean for the i-th chain is 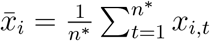. The within-chain variance is

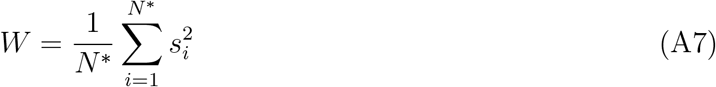

where the variance for the i-th chain is 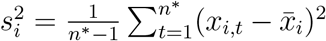. The variance estimator is then

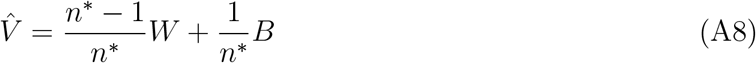

Finally, the Gelman–Rubin statistic is

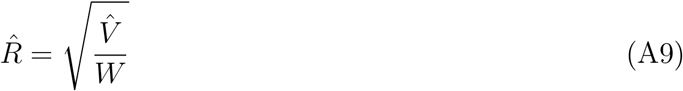

The stabilized samples of the clump coefficients will indicate that all chains are converging, then lead to 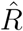 close to 1. In practice, we suggest splitting the samples into two halves (i.e., *N* ^*^ = 2) and using the heuristic threshold 1.01 or 1.02 that are commonly used in practice. We recommend to choose the number of stratum in the doubly-ranked method such that all the strata have satisfied a stabilization pattern of {*a*_*j,k*_, *b*_*j,k*_}_*k*=1,2,…,*K*_. All the strata in the simulation and real studies of the main text with coarsened exposures have the Gelman–Rubin statistic lower than 1.02. Figure S7 gives one example of the assessment.

**Supplementary Figure S7:**
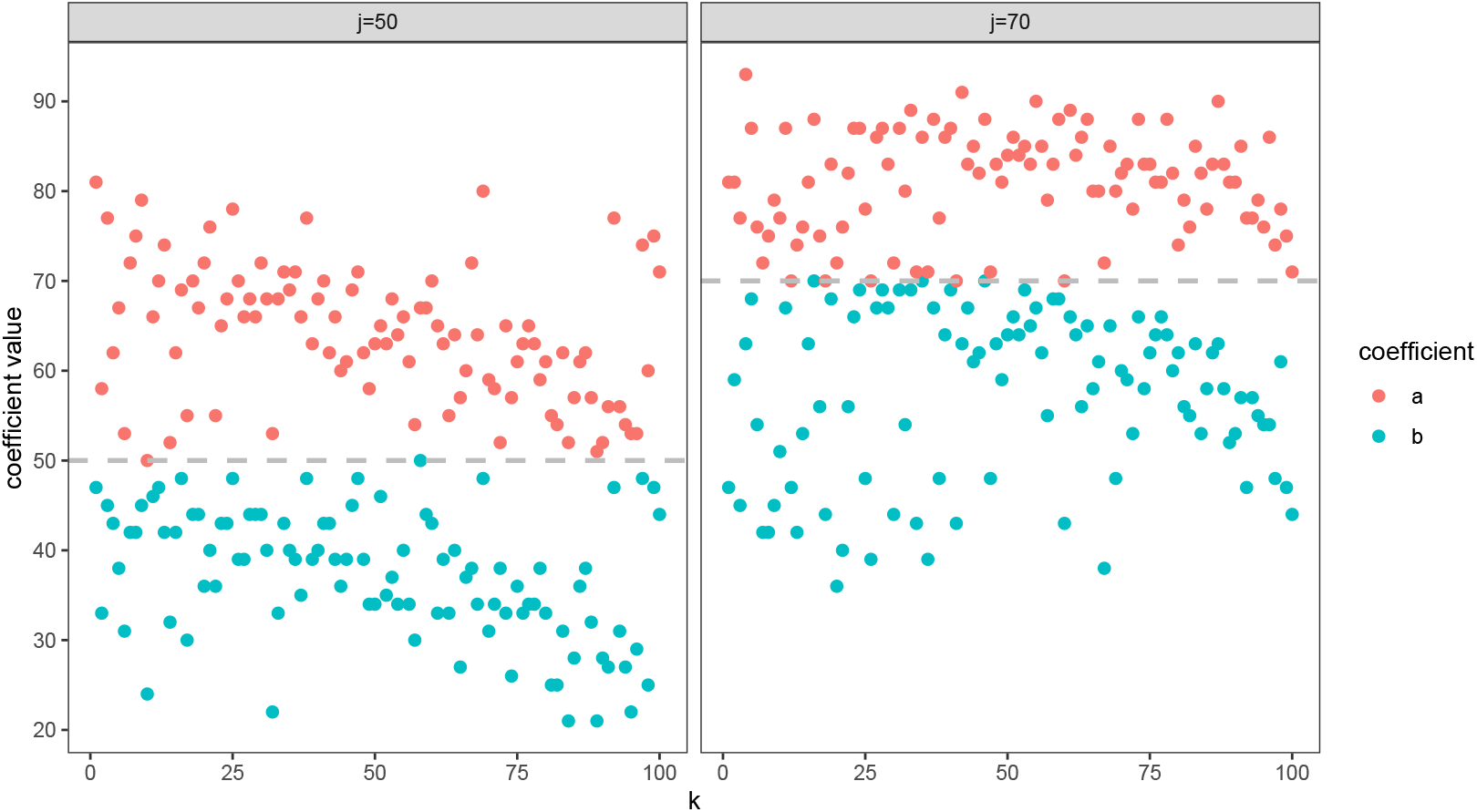
One example of the exchangeability assessment for the 50th and 70th stratum in a simulation with the doubly-ranked method. The exposure is coarsened and the sample size is 10000 in which 100 strata were created. The coefficients {*a*_*j,k*_, *b*_*j,k*}*k*=1,2,…,100_ are plotted against the stratum rank *k*. Left: the Gelman–Rubin statistics are 1.109 (for *a*_*j,k*_) and 1.150 (for *b*_*j,k*_) and the correlation of the confounder and the instrument is − 0.135 (strong evidence exchangeability is violated). Right: the Gelman–Rubin statistics are 0.999 (for *a*_*j,k*_) and 1.002 (for *b*_*j,k*_) and the correlation of the confounder and the instrument is 0.025 (no strong evidence exchangeability is violated).

## Text S2: Mathematical details for the causal effect of each stratum

We give the details of the target causal effect in each stratum used for the calculation of MSE and coverage rate in the main text. Assume the structural equation (for a stratum) is

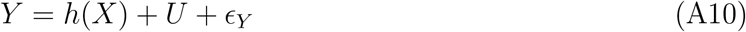

where *X, Y, U* and *ϵ*_*Y*_ are the exposure, the outcome, the unmeasured confounding and the exogenous variable respectively. We assume the effect shape *h*(·) is a differentiable function. Assume the IV assumptions (relevance, exchangeability, and exclusion restriction) for each stratum are satisfied. The MR will produce the estimator 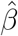 with the form

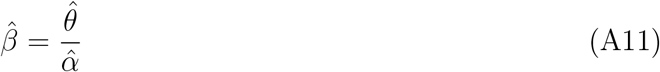

where 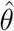 and 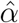 are obtained from the regression (in each stratum)

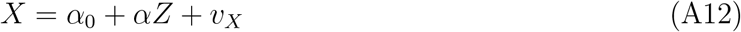

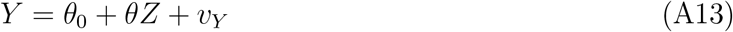

where *Z* is the instrument and *v*_*X*_, *v*_*Y*_ are the error terms. The MR estimator (A11) can also be derived with the same form by 2SLS and g-estimation. Let the exposure range is [*L, T*]. Now reconsider the equation (A10), which can be expressed as

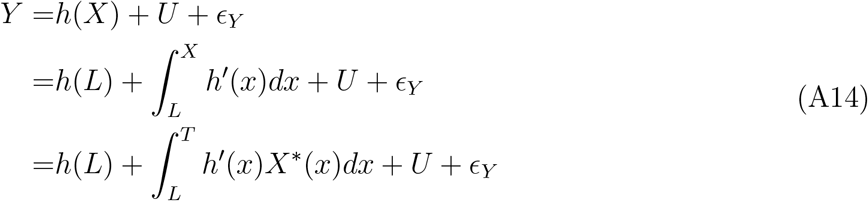

where *X*^*^(*x*) := *I* {*X* ⩾ *x*} is defined as the value-varying exposure. Consider the value-varying regression

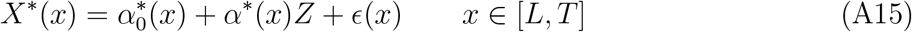

with 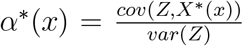 such that *cov*(*Z, ϵ*(*x*)) = 0 for any *x* ∈ [*L, T*]. The relevance condition ensures that *α*^*^(*x*) cannot always equal to 0 over [*L, T*]. Hence, we can further express (A14) as

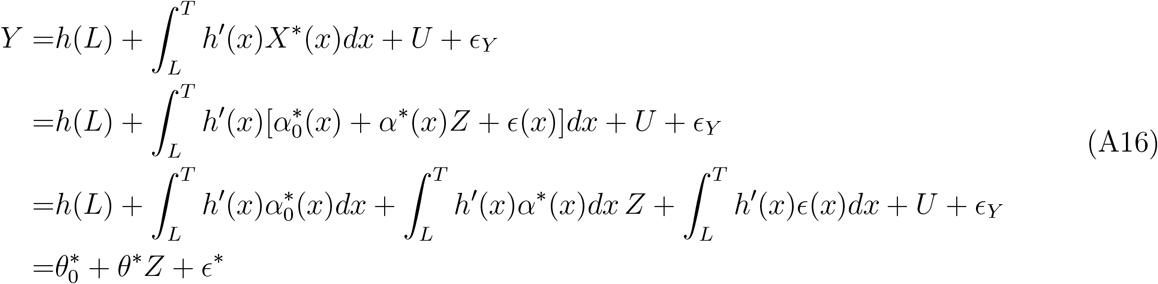

where

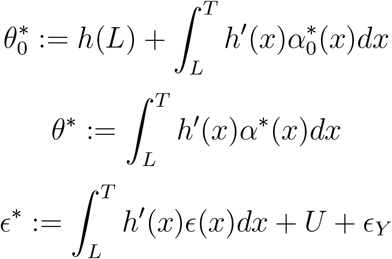

As 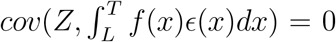 for general functions *f* (*x*) including *h*^′^(*x*) with regular conditions, *cov*(*Z, ϵ*^*^) = 0 due to the exchangeability condition. The slope estimator by fitting the regression (A13) will converge in probability as

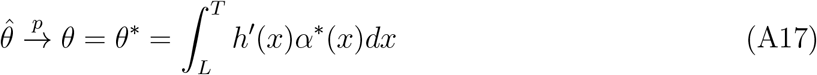

The MR estimator will then converge in probability as

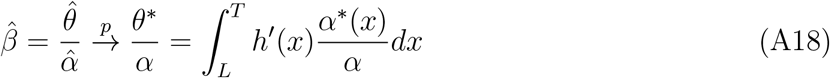

where the right-hand side is the target causal effect for the stratum. The weight function of the target effect over [*L, T*] is

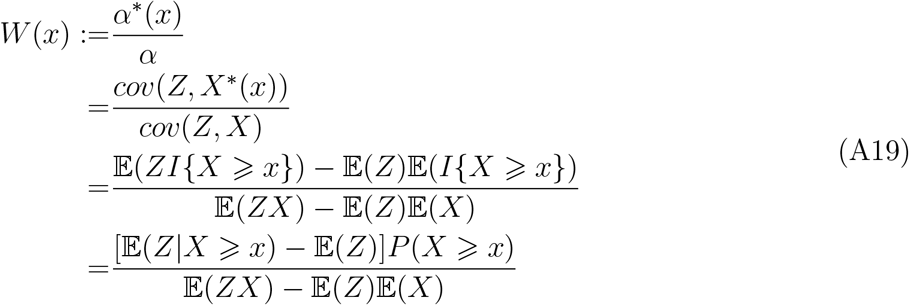

The weight function can be replaced by the empirical estimates or expressed via Monte Carlo simulation for each stratum. That is, we derive the estimate as

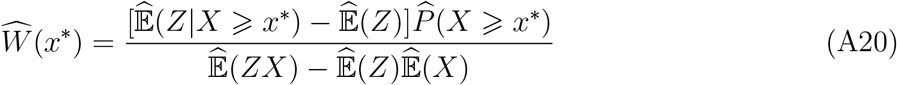

for multiple exposure candidate values *x*^*^ ∈ 𝒳. We then smooth 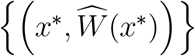 via smoothing methods like B-spline to obtain the estimated weight function 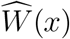.

## References

1. Davey Smith G, Ebrahim S. ‘Mendelian randomization’: can genetic epidemiology contribute to understanding environmental determinants of disease?. International Journal of Epidemiology 2003; 32(1): 1–22. doi: 10.1093/ije/dyg070

2. Burgess S, Thompson SG. Mendelian randomization: methods for causal inference using genetic variants. Chapman & Hall, Boca Raton, FL. 2021.

3. Burgess S, Davies NM, Thompson SG, EPIC-InterAct Consortium. Instrumental variable analysis with a nonlinear exposure–outcome relationship. Epidemiology 2014; 25(6): 877–885. doi: 10.1097/ede.0000000000000161

4. Staley JR, Burgess S. Semiparametric methods for estimation of a nonlinear exposure-outcome relationship using instrumental variables with application to Mendelian randomization. Genetic Epidemiology 2017; 41(4): 341–352. doi: 10.1002/gepi.22041

5. Amemiya T. The nonlinear two-stage least-squares estimator. Journal of Econometrics 1974; 2(2): 105–110. doi: 10.1016/0304-4076(74)90033-5

6. Newey WK, Powell JL. Instrumental variable estimation of nonparametric models. Econometrica 2003; 71(5): 1565–1578. doi: 10.1111/1468-0262.00459

7. Hall P, Horowitz JL. Nonparametric methods for inference in the presence of instrumental variables. The Annals of Statistics 2005; 33(6): 2904–2929. doi: 10.1214/009053605000000714

8. Horowitz JL. Applied nonparametric instrumental variables estimation. Econometrica 2011; 79(2): 347–394. doi: 10.3982/ecta8662

9. Mogstad M, Wiswall M. Linearity in instrumental variables estimation: Problems and solutions. tech. rep., Forschungsinstitut zur Zukunft der Arbeit. Bonn, Germany.; 2010.

10. Sun YQ, Burgess S, Staley JR, et al. Body mass index and all cause mortality in HUNT and UK Biobank studies: linear and non-linear mendelian randomisation analyses. British Medical Journal 2019; 364: l1042.

11. Malik R, Georgakis MK, Vujkovic M, et al. Relationship between blood pressure and incident cardiovascular disease: linear and nonlinear mendelian randomization analyses. Hypertension 2021; 77(6): 2004–2013.

12. Sofianopoulou E, others. Estimating dose–response relationships for vitamin D with coronary heart disease, stroke, and all-cause mortality: observational and Mendelian randomisation analyses. The Lancet Diabetes & Endocrinology 2021; 9(12): 837–846. doi: 10.1016/s2213-8587(21)00263-1

13. Cole SR, Platt RW, Schisterman EF, et al. Illustrating bias due to conditioning on a collider. International Journal of Epidemiology 2010; 39(2): 417–420. doi: 10.1093/ije/dyp334

14. Yusuf S, Wittes J, Probstfield J, Tyroler HA. Analysis and interpretation of treatment effects in subgroups of patients in randomized clinical trials. Jama 1991; 266(1): 93–98.

15. Small DS. Commentary: Interpretation and sensitivity analysis for the localized average causal effect curve. Epidemiology 2014; 25(6): 886–888.

16. Mason AM, Burgess S. Software Application Profile: SUMnlmr, an R package that facilitates flexible and reproducible non-linear Mendelian randomisation analyses. medRxiv 2021.

17. Greenland S. An introduction to instrumental variables for epidemiologists. International Journal of Epidemiology 2000; 29(4): 722–729. doi: 10.1093/ije/29.4.722

18. Martens E, Pestman W, Boer dA, Belitser S, Klungel O. Instrumental variables: application and limitations. Epidemiology 2006; 17(3): 260–267. doi: 10.1097/01.ede.0000215160.88317.cb

19. Hernán M, Robins J. Instruments for causal inference: an epidemiologist’s dream?. Epidemiology 2006; 17(4): 360–372. doi: 10.1097/01.ede.0000222409.00878.37

20. Swanson SA, Hernán MA. Commentary: how to report instrumental variable analyses (suggestions welcome). Epidemiology 2013; 24(3): 370–374. doi: 10.1097/ede.0b013e31828d0590

21. Lawlor D, Harbord R, Sterne J, Timpson N, Davey Smith G. Mendelian randomization: using genes as instruments for making causal inferences in epidemiology. Statistics in Medicine 2008; 27(8): 1133–1163. doi: 10.1002/sim.3034

22. Tudball MJ, Bowden J, Hughes RA, et al. Mendelian randomisation with coarsened exposures. Genetic epidemiology 2021; 45(3): 338–350.

23. Chen L, Davey Smith G, Harbord RM, Lewis SJ. Alcohol intake and blood pressure: a sys-tematic review implementing a Mendelian randomization approach. PLoS medicine 2008; 5(3): e52.

24. Cho Y, Shin SY, Won S, Relton CL, Davey Smith G, Shin MJ. Alcohol intake and cardiovascular risk factors: a Mendelian randomisation study. Scientific Reports 2015; 5(1): 1–13.

25. Larsson SC, Burgess S, Mason AM, Michaëlsson K. Alcohol consumption and cardiovascular disease: a Mendelian randomization study. Circulation: Genomic and Precision Medicine 2020; 13(3): e002814.

26. Sudlow C, Gallacher J, Allen N, et al. UK Biobank: an open access resource for identifying the causes of a wide range of complex diseases of middle and old age. PLOS Medicine 2015; 12(3): e1001779.

27. Astle WJ, Elding H, Jiang T, et al. The allelic landscape of human blood cell trait variation and links to common complex disease. Cell 2016; 167(5): 1415–1429.

28. Liu M, Jiang Y, Wedow R, et al. Association studies of up to 1.2 million individuals yield new insights into the genetic etiology of tobacco and alcohol use. Nature genetics 2019; 51(2): 237–244.

29. Burgess S, Davies NM, Thompson SG. Bias due to participant overlap in two-sample Mendelian randomization. Genetic Epidemiology 2016; 40(7): 597–608. doi:10.1002/gepi.21998

30. Wood AM, Kaptoge S, Butterworth AS, et al. Risk thresholds for alcohol consumption: combined analysis of individual-participant data for 599 912 current drinkers in 83 prospective studies. The Lancet 2018; 391(10129): 1513–1523.

31. Tasnim S, Tang C, Musini VM, Wright JM. Effect of alcohol on blood pressure. Cochrane Database of Systematic Reviews 2020(7).

32. Pajak A, Szafraniec K, Kubinova R, et al. Binge drinking and blood pressure: cross-sectional results of the HAPIEE study. PLoS One 2013; 8(6): e65856.

33. Maheswaran R, Gill JS, Davies P, Beevers DG. High blood pressure due to alcohol. A rapidly reversible effect.. Hypertension 1991; 17(6_pt_1): 787–792.

